# Multipath projection stereolithography (MPS) for 3D printing microfluidic devices

**DOI:** 10.1101/2024.07.18.604144

**Authors:** Zachary J. Geffert, Zheng Xiong, Jenna Grutzmacher, Maximilian Wilderman, Ali Mohammadi, Alex Filip, Zhen Li, Pranav Soman

## Abstract

Although many lab-on-chip applications require inch-sized devices with microscale feature resolution, achieving this via current 3D printing methods remains challenging due to inherent tradeoffs between print resolution, design complexity, and build sizes. Inspired by microscopes that can switch objectives to achieve multiscale imaging, we report a new optical printer coined as Multipath Projection Stereolithography (MPS) specifically designed for printing microfluidic devices. MPS is designed to switch between high-resolution (1×mode, ∼10µm) and low-resolution (3× mode, ∼30µm) optical paths to generate centimeter sized constructs (3cm × 6cm) with a feature resolution of ∼10µm. Illumination and projection systems were designed, resin formulations were optimized, and slicing software was integrated with hardware with the goal of ease of use. Using a test-case of micromixers, we show user-defined CAD models can be directly input to an automated slicing software to define printing of low-resolution features via the 3× mode with embedded microscale fins via 1× mode. A new computational model, validated using experimental results, was used to simulate various fin designs and experiments were conducted to verify simulated mixing efficiencies. New 3D out-of-plane micromixer designs were simulated and tested. To show broad applications of MPS, multi-chambered chips and microfluidic devices with microtraps were also printed. Overall, MPS can be a new fabrication tool to rapidly print a range of lab-on-chip applications.

## 1. Introduction

Microfluidic devices that enable the control and manipulation of microliter volumes of liquid are widely used for many applications. Photolithography remains the gold standard to make such devices, however due to the need for cleanroom and microfabrication facilities with technical expertise, specialized equipment, and labor-intensive steps (plasma bonding, PDMS molding, device assembly), this remains time and cost prohibitive especially for low production volume or high complexity designs. As an alternative to cleanrooms, 3D printing methods such as fused deposition modeling (FDM) and vat photo-polymerization (VPP) methods have been used to print microfluidic devices. However, due to low feature resolution of FDM (∼100μm), VPP methods have emerged as the method of choice to print high resolution microfluidic devices. VPP relies on light irradiation from a laser spot (vector scanning) or digital mask projections (DLP) to initiate polymerization and print 3D objects in a layer-by-layer manner. Since vector scanning approach is limited by long scanning times and complex process planning systems, DLP-VPP has emerged as the leading method for making microfluidic devices. In this method, a UV light source is spatially modulated by digital micromirror device (DMD) to generate pixelated light patterns derived from a sliced CAD model. DLP-VPP has been widely adopted to print miniaturized chips using custom and commercial printers and resins with channel sizes ranging from 25-150μm.[1-10] However, since pixel number is based on the number of micromirrors on a DMD chip (1920 × 1080), projection area is inversely proportional to the feature resolution. For instance, a build area of 48 mm × 36 mm[11] would have a resolution of ∼50µm while 1μm resolution can be only achieved by scaling down to a print area of only to 2 mm × 1 mm, an impractical size for most lab-on-chip applications.[12] To enable easy adoption by researchers, printed devices should fit onto a standard microscope slide (75mm x 26 mm), however this would require multiscale DLP-VPP strategies, as discussed below.[7, 13-19]

The most common type is the use of motorized step-stitching method that involves dividing the CAD file into a series of steps, moving either a motorized stage or the digital light engine by a defined distance before irradiation of pixelated images.[20, 21] Here, since print area per exposure does not change, feature resolution remains high. However, key limitations including – stitching errors between adjacent regions despite sophisticated image processing methods and, longer fabrication times due to multiple stage movement and exposure steps remain. To reduce print times, concurrent light projection and stage movement have also been developed,[22-24] however high image refresh rate to avoid motion blurring during printing requires a custom graphics hardware which has limited its utility in the field. The strategy of mounting multiple projectors to cover a larger area involves high costs and alignment issues. Another strategy is to use two distinct light sources, one to print low-resolution and typically internal features and second a high-resolution laser to print contours.[25-27] To maintain high print speeds and resolution, pixel blending methods have been developed however this remains computationally prohibitive.[28] Hybrid machines have also been built that integrate laser scanning using galvo-mirrors with DLP-VPP, however high cost, complex process planning, and low print speed and spot positioning errors during laser scanning remains a challenge.[29] Recently, two-axes galvo mirrors combined with a custom f-theta lens and novel hopping light DLP offers promising solutions for multiscale printing.[30] Combining vector scanning and DLP-VPP involve complex process controls to coordinate slicing algorithm with laser path planning and mask generation.[25, 31-33] Machines with integrated vertical and rotatory degree of freedom have also been used for large-area printing, however complex control systems have limited its utility in the field.[34] Overall the complexity of such multiscale DLP platforms prevent their adoptions within non-specified broader communities.

With ease of use as our inspiration and microfluidic chips as our target application, we set out to design a printer that could print devices that would fit onto a standard microscope slide (75mm × 26 mm) yet maintain a feature resolution of ∼10µm, without significantly increasing process complexity, printing time, or hardware costs. Here, we report a new multipath projection stereolithography (MPS) printer capable of rapid multiscale printing of parts as large as 30mm × 60mm (1.8 × 2.36 inch^2^) sized microfluidic devices and structures with ∼12µm resolution. MPS consists of a single light source and two optical configurations that can be switched between 3× mode (resolution ∼30µm) and 1× mode (resolution ∼12µm) to realize multiscale microfluidic devices. Both lateral and vertical resolution for each mode were characterized. The ability to print 3D structures with complex designs was demonstrated using an Empire State Building and alveolar model with complex internal fluidic topologies. Using micromixers as a test case, we show that MPS can rapidly design and print devices with variations in fin type based on target mixing efficiency derived from fluid flow simulations. We also show the printing and testing of micromixers with complex 3D out-of-plane channel topologies. Lastly, we report that MPS can be used to rapidly design and print microfluidic devices that cannot be printed by 1× or 3× mode used in isolation; here 3× mode was used to print macroscale features, while smaller microscale features were printing using 1× mode.

## 2. Materials and methods

### 2.1. PEGDA Prepolymer Preparation

Poly (ethylene glycol) diacrylate (PEGDA, Mn = 250) and the photo-initiator, phenylbis (2,4,6-trimethylbenzoyl) phosphine oxide (IRGACURE 819) were purchased from Sigma-Aldrich and used without any further modifications. The photo-absorber 2-Isopropylthioxanthone (ITX) was purchased from Tokyo Chemical Industry and used without further modifications. The prepolymer solution was composed of PEGDA (100% v/v) with IRGACURE 819 (1% w/v) and ITX (1.5% w/v). The prepolymer solution was mixed with a stainless-steel stirrer, then vortexed and placed in a water bath at 37°C repeatedly until IRGACURE 819 and ITX had dissolved.

### 2.2. Fabrication

The material vat consists of a polystyrene Falcon brand 100×15mm petri dish with a poly(dimethylsiloxane) (PDMS) buffer cured to the bottom of the dish. Approximately 3.5g of PDMS is poured into the petri dish, vacuum degassed to remove entrained air bubbles, and heat cured at 60°C overnight.

### 2.3. Methacrylation of Glass Coverslip

Glass coverslips were immersed into 10% (w/v) NaOH solution for 30 min, and washed in DI water, 75% (v/v) ethanol, and 100% ethanol (performed twice for 3 min for each wash). The coverslip was subsequently dried using nitrogen. The dried coverslips then underwent methacrylation by immersing them for 12hr in a solution comprised of 85 × 10^−3^M 3-(trimethoxysilyl) propyl methacrylate (TMSPM, Sigma) and ethanol solution with acetic acid (pH 4.5). Finally, the coverslips were washed with ethanol three times and baked for 1hr at 100 °C.

### 2.4. Fluorescent dyes

2 mg/mL 150 kDa FITC-dextran and 1 mg/mL 70 kDa Rhodamine-dextran were mixed in a water solution.

### 2.5. SEM

For obtaining the SEM (JSM 5600, JEOL, Japan) images, samples were separated from their printing mount, washed with ethanol, and dried. Then, samples were sputter coated (Vacuum Desk V, Denton, Moorestown, NJ) for 45 seconds with a layer of gold and imaged under the SEM with 10kV accelerating voltage.

### 2.6. Micro-CT Analysis

Following printing, the microfluidic chips were washed with ethanol and placed on a solid 3D printed base with double sided tape to prevent movement. The base was placed inside a 20mm diameter sample holder for micro-CT imaging (micro-CT 40, Scanco Medical AG, Brüttisellen, Switzerland). Imaging was conducted at a 10µm isotropic voxel resolution using 55kV, 145mA, and a 200ms integration time. Following scanning, the reconstructed images (.isq files) were transferred into Materialise Mimics, a 3D medical image segmentation software, for analysis. Images were then cropped to isolate the microfluidic chips, and a global threshold of 200 mg HA cm^-3^ was applied. A 3D reconstruction was generated from this data and exported as an STL file for visualization purposes.

### 2.7. CFD

To reduce the cost and time for physical prototypes and accelerate the microfluidics development process, we employed computational fluid dynamics (CFD) simulations implemented in ANSYS Fluent to allow rapid prototyping in a virtual environment. CFD studies provide us detailed insights into flow patterns, mixing features and concentration distributions in microfluidic channels with various microscale fins. This predictive capability helps in anticipating the behavior of the fluid flow and mixing before an actual microfluidic channel is fabricated. CFD simulations are used to optimize the design of microfluidic channels, including adjusting channel geometries and structures, flow rates, and other parameters to achieve efficient mixing along the flow. Details of the CFD model and simulation results and corresponding analysis are included in **SI5**.

### 2.8. Components/devices for system design

The optical and opto-mechanical components are purchased from Thorlabs, Edmund Optics and RPC photonics. Other customized mechanical components such as a rotator for engineered diffuser, polymer vat, Z stage etc. as well as several alignments assisted components are specifically designed and machined in-house or directly purchased from McMaster. Laser source was previously purchased from Toptica and LED light source was purchased from Golden-Scientific, while the DMD development kit (0.95’ UV 1080p) was previously purchased from DLi innovation.

### 2.9. Lens design/Mechanical design

The lens design and optical analysis are performed in ZEMAX software. The mechanical design is performed in Autodesk Inventor.

### 2.10. System control

Control software was developed using LabView (National Instruments).

### 2.11. Laser speckle characterization

The laser speckle pattern or illumination uniformity is characterized using beam profiler (Newport) at the plane of the polymer vat.

### 2.12. Characterization of absorption spectrum

Photo-initiators and photo-absorbers at 0.001% w/v were dissolved in PBS or ethanol, placed in 4.5mL plastic cuvette (Fisher Scientific), and then characterized using a UV-VIS spectrophotometer (Thermal Fisher) to measure their absorption spectrum from 300-800 nm.

### 2.13. Characterization of transparency

UV-VIS spectrophotometer (Thermal Fisher) was used to measure the transmission spectrum (400 - 800nm) using 4.5mL 100% PEGDA polymer mixed with candidate photo-absorbers (0.01%). DSLR camera was used to snap pictures to visualize the transparency.

## 3. Results and discussions

### 3.1. MPS system design

MPS is inspired from conventional microscopy which can switch between high- and low-resolution objectives to change the image size and feature resolution, allowing multiscale imaging. MPS utilizes two optical paths that can be switched as desired to achieve necessary print size while maintaining high resolution **(Fig. 1a)**. Using a test case of microfluidic devices, which fit on a standard cover slide, MPS is designed with two pathways named 1× and 3× to realize a maximum print area of 30mm × 60mm while maintaining the ability to print at a resolution of 12µm.

**Figure 1.**
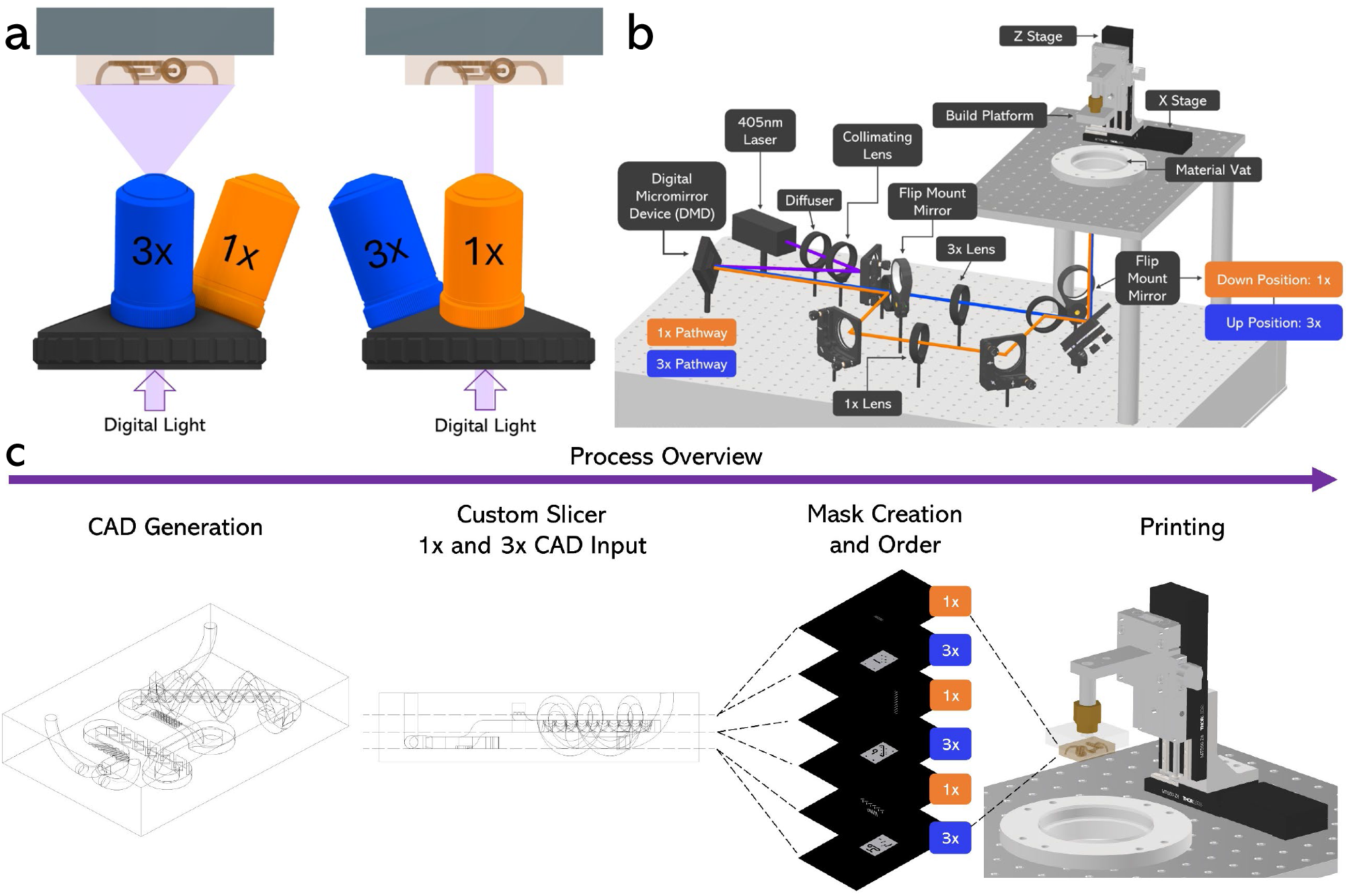
**a** Demonstration of MPS concept with two modes similar to that of multi-resolution microscopy. **b** Schematic of multipath projection stereolithography (MPS) printer setup. The flip mount mirrors allow for the system to switch between 1× and 3× quickly, directing the patterned light on two different pathways, 1× (Orange) and 3× (Blue). When the flip mount mirrors are in the down position, constructs are printed at a 1× scale and when the flip mount mirrors are in the upward position, constructs are printed at a 3× scale. **c** MPS process overview including CAD generation, 1× and 3× CAD input to custom slicer, mask creation and order, and finally printing.

First, we tested two illuminations systems (365 nm fiber coupled LED, 405 nm semiconductor laser) with our multi-projection system. However, LED was not selected due to the low transmission efficiency (<50%) lens used in the setup, and associated challenges related to energy efficiency and influence of NA (0.5) on the projection lens. Details are explained in SI. Therefore, the 405nm laser was selected as the light source in our setup. Briefly, the laser was collimated using a plane-convex lens with focusing length 150 mm (Thorlabs), and an engineered diffuser (RPC photonics Inc., USA) was used to convert the Gaussian profile of the laser beam into Top-hat profile (**Fig. S1**). This was done to obtain uniform illumination intensity before projecting onto the digital micromirror device (DMD). The lens selection was based on the divergence angle of the engineered diffuser and the illumination area of DMD (25.4mm). During our testing, we found that laser speckle, a common problem due to coherence property of the laser, negatively affects the illumination uniformity. This issue was solved by designing and building a setup to rotate the diffuser and obtain an illumination uniformity greater than 85%.

Here the system resolution, the smallest distance between two features, is largely designed based on the DMD micromirror size (∼10µm, with a gap of 1µm between mirrors). We chose a smaller numerical aperture (NA, 0.04) in both the illumination and projection setups to have sufficient depth of focus and minimize errors in opto-mechanical alignments between the Z stage and the bottom of the vat. Precision stages can resolve this issue, however our choice of having the depth of focus over 200µm was motivated by lowering costs and reducing the complexity. In our case, the maximum distortion across the field of view@12 mm is less than one pixel of DMD (10.8 µm), which is less than 0.1%. For two adjacent pixels in DMD to be resolved at the image plane, the modulation transfer function or MTF@ 50lp should be more than 0.5 for 1× mode while MTF@20lp should be more than 0.5 for 3× mode. System analysis, as performed by ZEMAX, showed that both modes reach their diffraction limits. SI shows specifics about the 2D layout of both optical systems, optimized using three main field of view (FOV=0, 0.707, 1). Results show that (i) RMS spot radius at all FOVs is less than airy radius (18.41µm) showing that the system performance reached its diffraction limits, (ii) maximum distortion across field of views is less than 0.1%, (iii) MTF at all FOVs remains under the diffraction limit; for 1× mode, MTF at full field of view remains less than 0.5. **Figures S1-3** provide additional specifics about the laser illumination system.

Thus, the final setup consists of a DMD, an engineered diffuser, a 405nm CW laser, lenses for 1× and 3× modes, two flip mount mirrors, and an X and Z stages **(Fig. 1b)**. Light irradiated from the laser was diffused, creating a uniform intensity distribution, and further collimated by illumination optics before directing onto the DMD. The DMD used in this system consists of 1920×1080 array of micromirrors with single pixel resolution of 10.8 µm. Following the DMD, the laser path can be directed onto two different pathways, 1× (Orange) and 3× (Blue) based on the position of two flip mount mirrors (Down:1×, Up:3×). Both pathways are directed upwards by a 45-degree mirror towards the material vat, where material can be placed for printing. On the build platform there is an X and Z stage to allow layer by layer printing in the Z and movement of 1× features in the X direction. MPS uses a simple process, beginning with a CAD generation followed by a custom MATLAB 3D slicer which provides masks output in the correct ordering between 1× and 3× **(Fig. 1c)**.

### 3.2. Automation of MPS

To minimize alignment errors and increase repeatability, we automated both design and printing aspects of MPS. This automation process is explained in **Fig. S4**. Briefly, 3D CAD models, designed using Autodesk Inventor, containing macroscale and microscale features to be printed via 3× and 1× modes of MPS respectively. A custom 3D slicer, developed in MATLAB, was used to generate image files for both modes. The process flow starts with user selection of CAD files to be printed with 1× and 3× modes, followed by choosing the layer heights for each mode, and the layer number where the modes will be switched from 3× to 1× mode with flip mount mirror positions for each mode. Since design tradeoffs make perfect alignment between modes challenging, simple image processing, such as adding an image offset, can be used to compensate for any alignment errors. MPS can also be operated in individual 3× or 1× modes. A graphical user interface (GUI) is used to monitor, and control various aspects of MPS such as stage position, DMD parameters, print duration, layer heights for each mode, detachment distance, display mask images, mirror position and other things. Before printing, stage is lowered in a resin filled PDMS vat to set the start position, and other parameters related to laser power, image files, stage, and number of layers. A step-by-step process flow, and associated control algorithm files are provided in the **SI3** section.

### 3.3. Characterization of MPS

Resin selection remains a difficult challenge for any additive manufacturing project. In this work, we were most focused on rapid iteration of microfluidic devices, so a reliable workhorse material was required. Resin optimization was performed with this in mind, and the formulation was identified to include PEGDA (250 Da) as the base material, Irgacure 819 as the photo-initiator, and ITX as the photo-absorber **(Fig. S5)**. This formulation was used in the remainder of the work. To characterize the resolution of MPS, the lateral-resolution was examined first. Digital masks of line patterns designed with pixel numbers varying from 1-32 pixels were printed using both 1× and 3× modes individually **(Fig. 2a)**. and a digital microscope (HIROX, Japan) was used to measure the linewidths, showing XY-resolutions for 1× and 3× optical paths to be 12.93±1.32µm and 30.13±2.09µm respectively, giving an actual magnification ratio of approximately 2.8 **(Fig. 2b)**. Z-resolution was examined next by controlling the exposure dose, a function of light intensity and time. A ladder structure was printed with varying exposure times while maintaining a constant exposure intensity of 3.5mW/cm^2^ **(Fig. 2c)**. Results were fitted using the Beer-Lambert Equation, showing Z-resolution variation between 12.68 to 132.75µm for an exposure time-range of 0.3 to 3 seconds **(Fig. 2d)**. Based on these results, we choose a layer height of 50µm using an exposure time of 0.8 seconds per layer.

**Figure 2.**
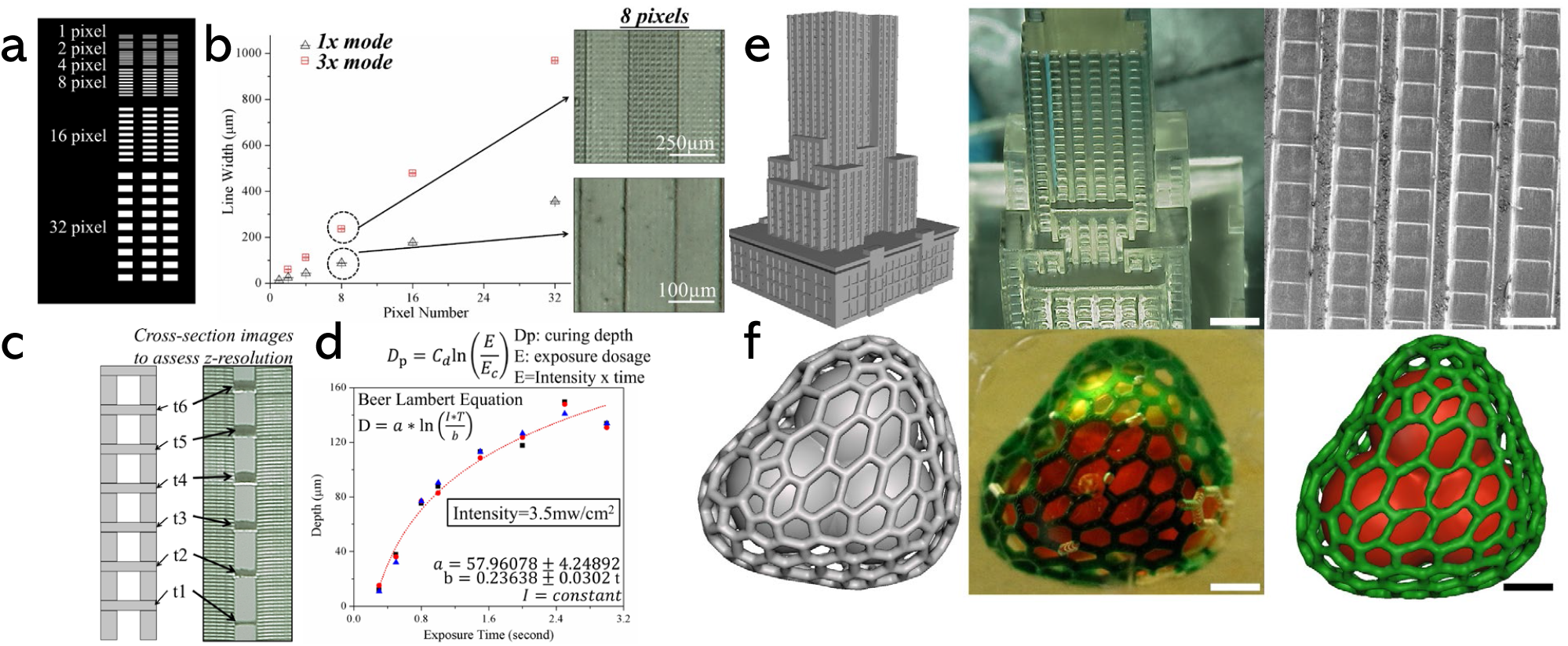
**a** Line pattern design varying from 1-32 pixels for XY-resolution characterization. **b** Measured linewidths plotted vs pixel number show 1× resolution of 12.93±1.32µm and 3× resolution of 30.13±2.09µm. **c** Z-resolution characterization using ladder structure completed by varying exposure time while maintaining constant light intensity. **d** Ladder results fitted using Beer-Lambert Equation. **e** Empire State Building CAD, printed structure imaged under HIROX, and SEM image. Scale bars are 2mm and 500µm, respectively. **f** Alveoli model including two individual interconnected structures CAD, printed alveoli structure imaged under HIROX, and microCT reconstruction of construct. Scale bars are 1mm.

The capability of printing complex designs was tested using 3× mode of MPS. First, the Empire State Building was modeled **(Fig. 2e)** and printed. Images taken using HIROX distinguish microscale windows of building and Scanning Electron Microscopy (SEM, JEOL5600, Japan) images distinguish features printed using single pixel light exposure. With an approximate volume of 1.4cm^3^, this structure was printed in less than 5 minutes. Second, an alveoli-mimicking structure, with complex hollow topologies, was tested. A representative alveoli found in the human lungs was designed **(Fig. 2f)** and printed. It consisted of two independent hollow features, including an interconnected air sac (red food dye) and a network of microchannels surrounding the air sac structure representing blood capillaries (green food dye). To accurately characterize the printed construct, micro computed tomography (micro-CT 40, Scanco Medical AG, Brüttisellen, Switzerland) was performed. Results show high printing accuracy.

### 3.4. Rapid prototyping of microfluidic mixers

Microfluidic mixers were chosen as a test-case to demonstrate MPS’s capability printing microscale features in any defined location within a macroscale device. With insight from the literature, three microfluidic mixer devices were designed and printed using MPS. In the field, the three most common mixers utilized included 3D spiral fins forcing fluid horizontally and vertically, fixed solid wall fins where fluid is forced through a pathway, and a herringbone pattern where fluid flows over the top.[35-38] Additionally, most mixers followed a serpentine pattern to maximize channel length and mixing efficiency within in fixed chip size. With these specifics in mind, the first mixer was designed with a 500µm wide serpentine channel within a 12×8mm microfluidic chip. This included two inlets and one outlet. The channels were 400µm in height and this design was drawn with 100µm fixed solid fin walls at 45deg. This allowed for a 300µm opening for fluid flow between the fins. The overall chip and serpentine were printed in under 10 minutes using 3× mode and the fixed solid fin walls were printed using 1× mode of MPS **(Fig. 3a)**. Top view of CAD model is shown (**Fig. 3b)**. We included computational modeling and experimental results to have a complete approach, tunable for different applications. For this first mixer, experimental analysis was performed first to validate custom computational fluid dynamics (CFD) algorithm data to develop a predictable model. CFD analysis was performed using ANSYS. To certify the laminar flow conditions, we conducted a flow simulation (without diffusion) for each case to calculate the maximum velocity and the corresponding Reynolds number in the channel. **(SI5)** To assess mixing efficiency experimentally, fluorescent dyes were chosen.[38] 150 kDa FITC-dextran and 70 kDa Rhodamine-dextran were flowed in each inlet at 5 µL/min, controlled by a syringe pump. Fluorescent images were acquired throughout the chip; the inlet was chosen as the baseline for mixing efficiency **(Fig. 3c)**. The mixing ratio was calculated in MATLAB by computing the percent overlap between the normalized fluorescence intensity profiles at the start and end of the flow channel.[38] Image of end point is shown with mixing ratio graphs **(Fig. 3d)**. The final mixing ratio of this mixer was determined to be 83.25% **(Fig. 3e)**. CFD results of the top view of the channel, and cross sections of the start and end sections of the channel are shown **(Fig. 3f)**. Mixing efficiency was determined to be 83.39% using methods described in **Figures S6-9**. Additional images including an isometric CAD view, no roof internal view of fin design, fluorescent image from middle section, and SEM characterization can be seen in **Figure S16a**. Experimental results align well with the CFD mixing efficiency results, further emphasizing the print quality and success of the MPS system for this application.

**Figure 3.**
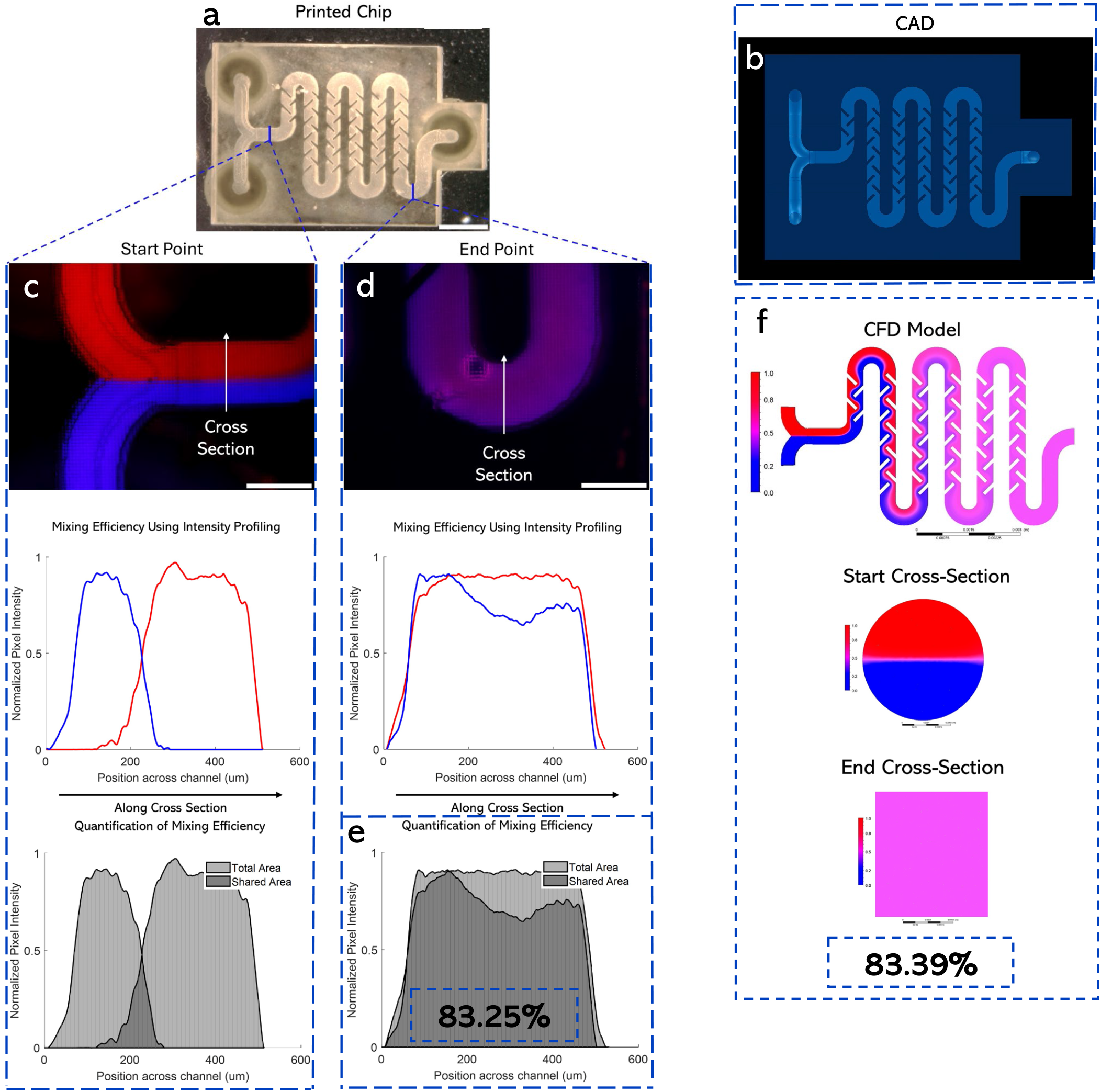
Microfluidic mixers as an example of rapid iteration of printable microfluidic devices. **a** Fixed solid wall printed result top view. Scale bar is 2.5mm. **b** Top view of fixed solid wall CAD model. **c** Fluorescent image of start position and mixing efficiency graphs using intensity profiling. Scale bar is 500µm. **d** Fluorescent image of end position with mixing efficiency graphs. Scale bar is 500µm. **e** Quantification of mixing efficiency using normalized pixel intensity across channel position. **f** CFD results including top view and start/end cross section views.

For validation of our rapidly iterative printing approach, two additional microfluidic mixers were designed with the same overall chip size, serpentine channel width and height, but the mixing feature to be printed by 1× was changed. CAD design to printed structure can be completed in under 2 hours. The second mixer was designed with 100µm wide 3D spiral fins, using a 1440deg twist, creating 8 rotations per straight section of the serpentine **(Fig. 4a)**. Here, the CFD model, validated using the fixed solid design, was used to calculate mixing efficiency before printing the device with MPS **(Fig. 4b)**. Mixing efficiency was determined to be 99.18% from CFD analysis using methods illustrated in **Figures S10-12**. Again, the chip was printed with 3× and the 3D spiral fins with 1× **(Fig 4c)**. Fluorescent images and mixing efficiency are illustrated **(Fig. 4d-e)**. The final mixing ratio of the second mixing design was determined to be 90.55%. Additional images including an isometric CAD view, no roof internal view of spiral section, fluorescent image from middle section, and SEM characterization can be seen in **Figure S16b**. Experimental and computational results of mixing efficiency match well.

**Figure 4.**
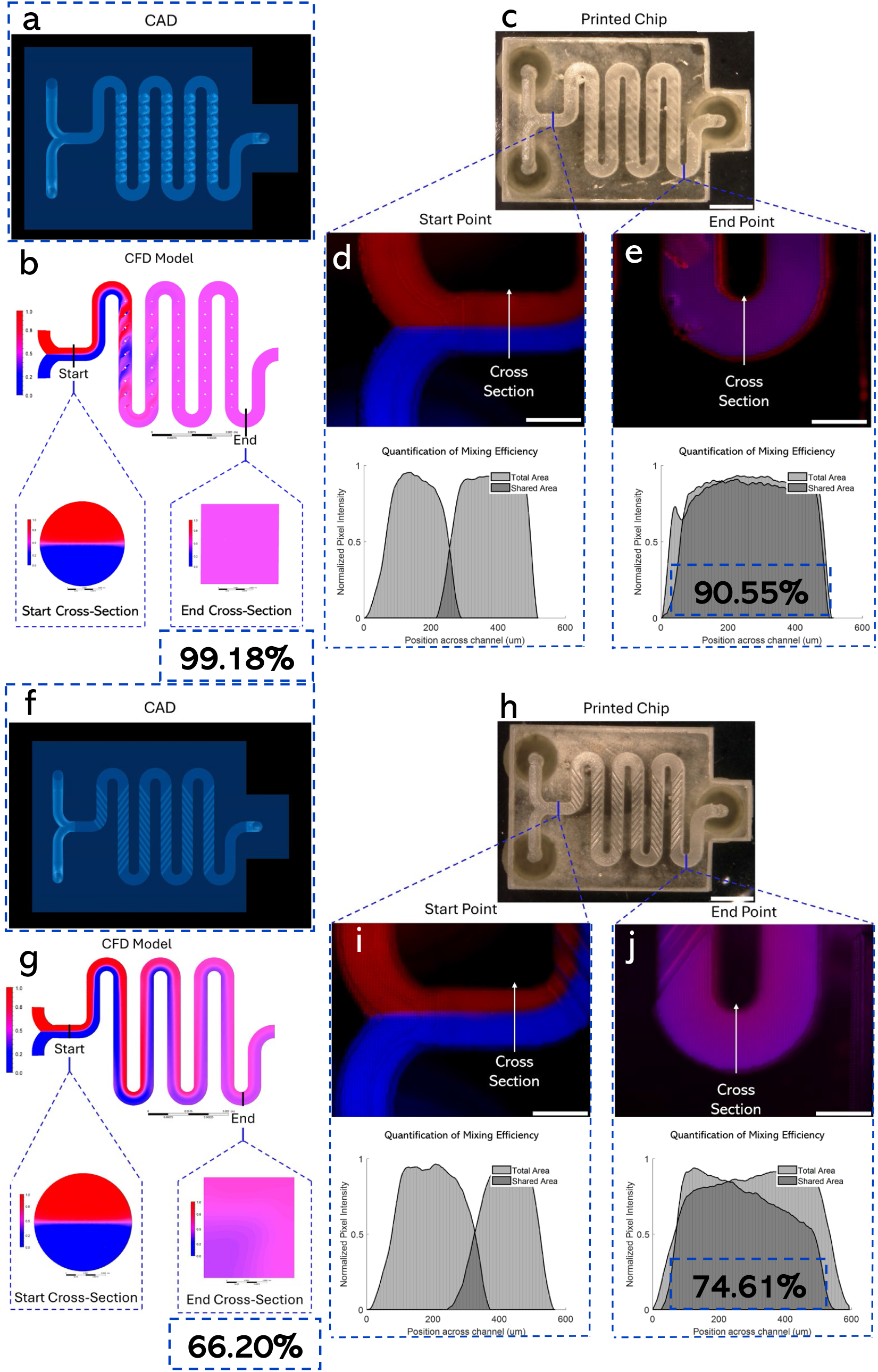
**a** Top view of 3D spiral CAD model. **b** CFD results including top view and start/end cross section views. **c** 3D spiral printed result top view. Scale bar is 2.5mm. **d** Fluorescent image of start position and mixing efficiency graph using intensity profiling. Scale bar is 500µm. **e** Fluorescent image of end position with mixing efficiency graph illustrating quantification of mixing efficiency using normalized pixel intensity across channel position. Scale bar is 500µm. **f** Top view of herringbone CAD model. **g** CFD results including top view and start/end cross section views. **h** Herringbone printed result top view. Scale bar is 2.5mm. **i** Fluorescent image of start position and mixing efficiency graph using intensity profiling. Scale bar is 500µm. **j** Fluorescent image of end position with mixing efficiency graph. Scale bar is 500µm.

The third mixer was also designed with the same chip size, and serpentine characteristics, and included a herringbone pattern on the bottom of the channels. The serpentine pattern was printed with a 100µm height, 100µm gap between each fin, and an experimentally determined 35.6deg angle **(Fig. 4f)**.[35] CFD analysis determined mixing efficiency to be 66.20% **(Fig. 4g)** using methods illustrated in **Figures S13-15**. Printed chip result is shown by a top view **(Fig. 4h)**. Top view fluorescent images from the start and end points are shown along with mixing efficiency graphs **(Fig. 4i)**. For the herringbone fin design, mixing efficiency was shown experimentally to be 74.61%. Similar images including an isometric CAD view, no roof internal view of herringbone pattern lining bottom of channels, fluorescent top view image, and SEM characterization can be seen in **Figure S16c**. Overall, experimental results from all three fin designs showed consistency with CFD mixing efficiency results.

### 3.5. Complex 3D microfluidic mixers

Using inspiration from the features designed in the previous mixers, two complex 3D mixers were designed to further highlight the unique capabilities of MPS, particularly its ability to print complex structures in multiple locations in 3D. In the first CAD, channels were designed to flow and overlap on three planes. On each plane, a different mixing feature design was incorporated. The first utilizes a herringbone design and fixed solid fin wall structure. The second plane and third planes feature a 3D spiral design inside the channel and an array of microdots, respectively. Again, the top and side view of the 3D CAD is shown **(Fig. 5a)**. The channels overlap each other multiple planes including a spiral around a section of the channel. CFD highlighted the efficiency of the design at 93.10% **(Fig 5b)**. Furthermore, microCT was performed for printing validation, and results are shown **(Fig. 5c)**. Finally, the printed result is shown from a top view with a final mixing efficiency of 91.01% **(Fig 5d)**.

**Figure 5.**
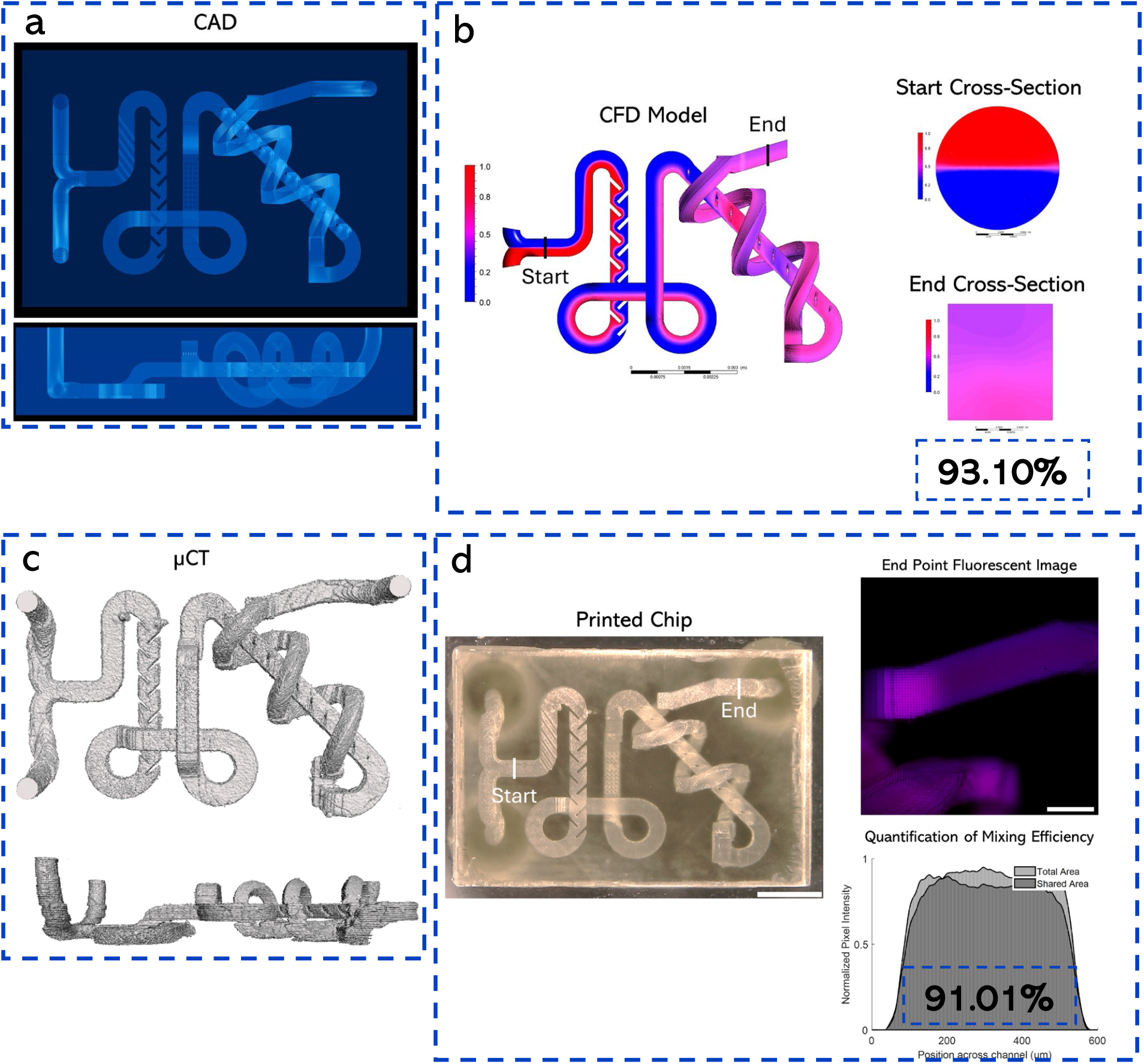
**a** CAD of complex 3D microfluidic mixer including top and side view. Features on three planes include herringbone, fixed solid fin wall, micro dots, and 3D spiral. **b** Printed result, top view. **c** microCT reconstruction of microfluidic mixer with same model views. **d-e** CFD results include top view and start/end cross section views. Scale bar is 2.5mm.

In the second complex CAD design, channels were designed to flow and overlap on two planes. On the bottom of channels throughout the chip, the same herringbone pattern was used as the mixing design. The top and side view of the 3D CAD is shown **(Fig. S17a)**. CFD analysis was performed pre-printing to allow for further optimization of the design. End cross-section result is shown, highlighting the high efficiency of the design at 98.31% **(Fig. S17b)**. To accurately characterize the printed mixers, we used microCT. Results shown in the same views exhibit excellent mimicry of the original CAD design **(Fig. S17c)**. The printed result is shown from a top view and fluorescent mixing efficiency is highlighted at 90.18% **(Fig. S17d)**. A summary of all chip mixing efficiencies is shown in **Figure S18**.

### 3.6. Other microfluidic devices using MPS

To demonstrate the feasibility of using MPS to print large-scale devices with high resolution, we printed a simple cell trapping microfluidic device using both 1× and 3× modes. The microfluidic chip base, 40mm × 20mm, was printed using 3× mode with three microtrap arrays embedded within the device printed using the 1× mode **(Fig. 6a)**. The height of the channel was 500µm, while the height of the microtraps was 100µm. Post-printing, a fluorescent microparticle solution (diameter=18.67µm, 10% v/v) was perfused with the central channel and a fluorescence microscope (Nikon, Japan) was used to capture images **(Fig. 6b-c)**. Printed microtraps, with a width of 60µm and a trapping opening of ∼20-22µm were able to trap single microparticles. Some of the traps were also able to trap more than one microparticles. These results demonstrate the potential of such a multiscale printer to rapidly print microfluidic devices for a range of applications.

**Figure 6.**
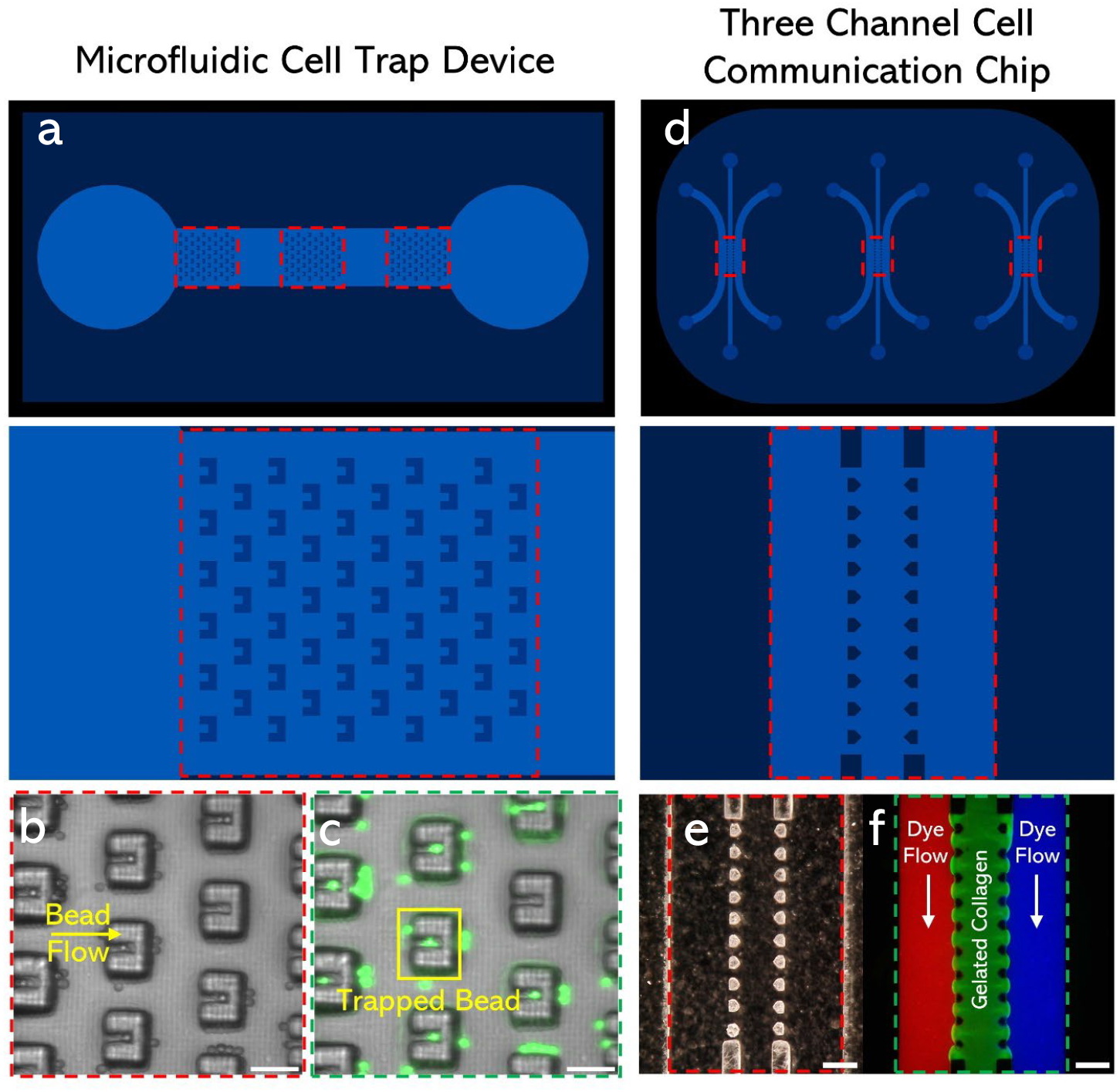
**a** CAD of large-scale microfluidic cell trap device. Three sets of cell trap arrays are printed by 1× on top of a 3× printed base. **b** Brightfield image of microparticle solution flowing through chip, microparticles are being stopped by traps. Scale bar is 100µm. **c** Fluorescent image of microparticles seen in traps. Scale bar is 100µm. **d** Three-channel cell communication microfluidic chip in triplicate (high-throughput). Scale bar is 2mm. **e** HIROX image of micro-posts created by 1× in between the three channels. Scale bar is 300µm. f Central channel filled with fluorescent collagen, and outside channels filled with microparticle solution. Scale bar is 300µm.

A second microfluidic device, a three-channel chip commonly used in 3D cell culture and organ-on-chip applications, was designed and printed using MPS. Three sets of the channel designs were printed within a single large-scale chip **(Fig. S19)** using the 3× mode, **(Fig. 6d)**, while micro-post arrays were printed using the 1× mode of MPS **(Fig. 6e)**. In a typical application, extracellular matrix or hydrogel solution perfused within the central chamber does not leak into the side channels. To demonstrate this, a 2% gelatin and 5% 2000kDa FITC-dextran solution was flowed into the central channel and allowed to thermally crosslink and solidify, before perfusing another fluorescent solution (150 kDa FITC-dextran) into the side channels **(Fig. 6f)**. Fluorescent microscopy image demonstrates no fluidic leakage between the three channels. These results highlight the unique ability of MPS to create high resolution microstructures in any location within a macro scale printed construct.

## 4. Conclusions

This study reported an alternative approach to fabricating multiscale microfluidic devices by combining a high-resolution and low-resolution mode into a single printing system. It overcomes certain tradeoffs found in the field between printing resolution and printing area. Conventional 3D printing methods, FDM for instance, have the ability to create large scale devices, however, lateral-resolution is limited to ∼100µm which is not sufficient for high quality microfluidic devices. Researchers have turned to VPP as a promising alternative, specifically DLP-VPP where higher resolution (<50µm) has been extensively reported. The major limitation of DLP-VPP is that its projection (build) area is inversely proportional to its feature resolution, which limits the creation of larger scale devices with high feature resolution. MPS utilizes DLP-VPP with inspiration from microscopy with multiple quick-change magnifications to overcome the aforementioned limitations. MPS does not come without its own limitations, but future work gives a promising path to address them. MPS is a powerful technology that can be applied to scales even larger or more importantly, smaller. With its concept demonstrated with 1× and 3×, there is an extendable capability for the system to be built with additional pathways including a 0.1× for even higher resolution features and/or 6× for larger scale devices. Though the existing MPS system utilized a single optimized material for microfluidic devices, as a DLP platform, it is inherently compatible with a diverse range of photo-crosslinkable materials which further extend its breadth of potential applications. One of the most challenging aspects of MPS is the physical alignment of the multiple pathways. It is difficult to perform attain alignment of any optical system, so an imaging processing algorithm was added within the slicer to mitigate misalignment. Utilization of different motorized optical components will be done in the future to minimize the need for such corrections. Finally, the slicer software that was developed relies on some user input to specify which features within the CAD model are to be printed with 1× and 3×. The existing algorithms can be augmented with more complex and intelligent detection abilities in the future. Such improvements to the print process will allow for our MPS platform to fabricate even more advanced structures for a wider variety of biomedical applications.

## CRediT authorship contribution statement

**Zachary J. Geffert:** Writing - review & editing, Writing - original draft, Data curation, Formal analysis, Investigation, Methodology, Software, Validation, Visualization. **Zheng Xiong:** Writing - review & editing, Writing - original draft, Conceptualization, Data curation, Funding acquisition, Methodology, Validation, Visualization. **Jenna Grutzmacher:** Writing - review & editing, Data curation, Visualization. **Maximilian Wilderman:** Data curation, Visualization. **Ali Mohammadi:** Writing - review & editing, Data curation, Formal analysis, Software, Visualization. **Alex Filip:** Conceptualization, Validation, Visualization. **Zhen Li:** Funding acquisition, Methodology, Project administration, Resources, Supervision. **Pranav Soman:** Writing - review & editing, Writing - original draft, Conceptualization, Funding acquisition, Methodology, Project administration, Resources, Supervision.

## Acknowledgements

We thank J. Horton (SUNY Upstate Medical University) for providing us with access to microCT. We also thank P. Kunwar and A. Poudel for their help in early system development and maintenance. Finally, we thank D. Fougnier for protocol review and A. Ram for his help with image processing.

Financial support for this project was provided by the National Institutes of Health (R21 GM141573-01) and National Science Foundation (SBIR Phase I to 3D Microfluidics LLC, Zheng Xiong, 2013942).

## Supplemental Information

### 1. Laser illumination system

A semiconductor continuous-wave laser (405nm with 8nm bandwidth, TOPTICA, Germany) is collimated using a plane-convex lens with focusing length 150 mm (Thorlabs). An engineered diffuser (RPC photonics Inc., USA) is used to convert the Gaussian profile of the laser beam into Top-hat profile (**Fig. S1**). This is important to obtain uniform illumination intensity before projecting onto the DMD. The lens selection was based on the divergence angle of the engineered diffuser and the illumination area of DMD (25.4mm). The focusing length of collimation lens is given by

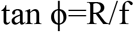

where ϕ is the divergence angle of the engineered diffuser (which is 5°), R is 0.5D the radius of lens aperture *(which should be larger than 12mm or ½ the size of DMD)*, and f is the focusing length of the collimation lens.

#### Problems encountered

Laser speckle, a common problem due to coherence property of the laser, negatively affects the illumination uniformity. This issue was solved by designing and building a setup to rotate the diffuser and obtain an illumination uniformity greater than 85%. To achieve high energy efficiency, the NA of projection system should be equal to or greater than the NA of the illumination system. However, higher NA will induce small depth of focus, which contributes to greater opto-mechanical misalignment. To solve this challenge, we choose laser illumination system with a smaller NA and thus decrease the strict requirement of NA for the projection lenses used in the system.

**Figure S1.**
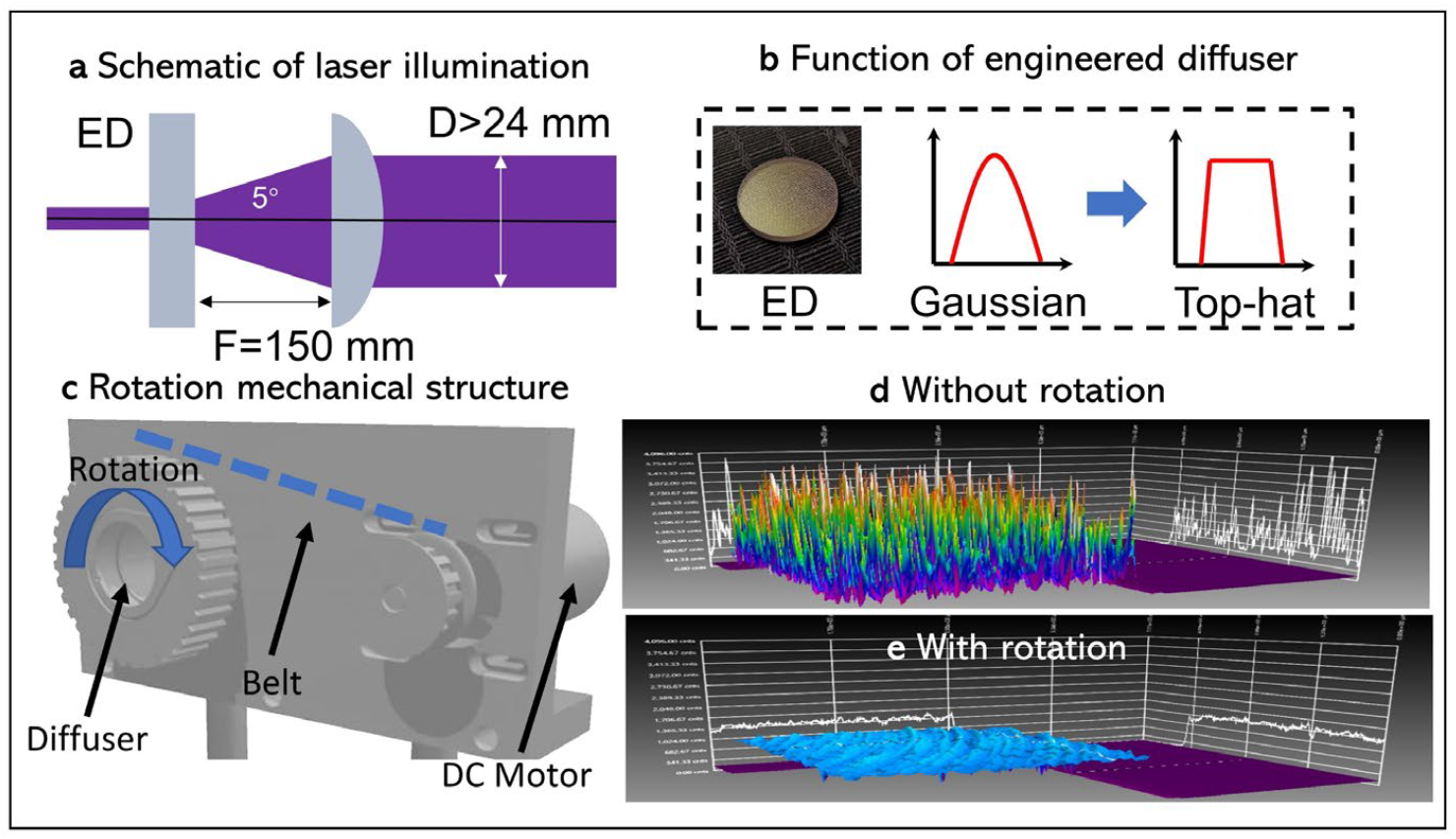
Laser illumination system. **a** Schematic of optical setup. **b** Function of engineered diffuser converting Gaussian profile of laser into Top-hat profile. **c** Rotating mechanical structure to generate uniform laser speckle. **d** Laser speckle profile with and without rotation.

### 2. Multi-path projection systems (1x and 3x) used in MPS

We have designed a projection system with two optical paths, 1x mode and 3x mode. 1x mode has a print area of ∼10mm x 20mm with a print resolution of **∼12µm** while 3x mode has a print area of 30mm x 60mm with a print resolution of **∼32µm**. Zemax was used to design the projection lenses based on several specifications, as discussed below.

#### Numerical aperture

NA of the projection system determines the resolution and the depth of the focus based on the following equations:

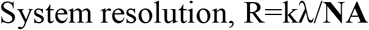

Where R is the system resolution, k is the lithography parameter, here, we defined as 0.5, NA is numerical aperture.

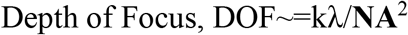

where NA=nsin ϕ, DOF is the depth of focus, λ is the wavelength of the light source, NA is the numerical aperture of the system, ϕ is the divergence angle of the engineered diffuser.

##### (a) System Resolution

Digital Micromirror Device (DMD) used in the setup consists of 1920×1080 micromirror array; size of each micromirror is 10µm with inter-mirror gap of 1µm. The resolution of the projection system should be low enough to not resolve the inter-mirror gap but high enough to resolve single micromirror (pixels), i.e, 10µm>R>1µm.

##### (b) Depth of focus (DOF)

DOF is the axial depth of the space on both sides of the image plane within which the image appears acceptably sharp. The stage, required to print large sizes, was found to be difficult to align with bottom surface of the vat; even microscale tilts of the stage resulted in opto-mechanical misalignment of ∼100µm **(Fig.S2)**. Precision stages and components, needed to address this issue, were found to be <u>prohibitively expensive</u>. As a result, a DOF of 200µm was finalized for this design. Based on this, NA will be 0.04. According to our system design, NA=nsinϕ=0.04 corresponds to divergence angle ϕ=2.3°. This divergence angle requires a high collimation; this is another reason why we choose the laser source in the illumination system as compared to LED.

**Distortion** describes the magnification in the image plane changes across the whole field of view. Since the maximum distortion at full field of view@12mm should be less than one pixel of DMD (10.8µm), the distortion at full-field of view should be less than 0.1%.

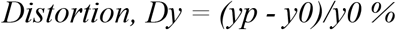

where Dy is the distortion, yp is the actual image height (in our case, it is ∼12mm±10.8µm, y0 is the ideal image height *(in our case, it is 12 mm)*.

**Figure S2.**
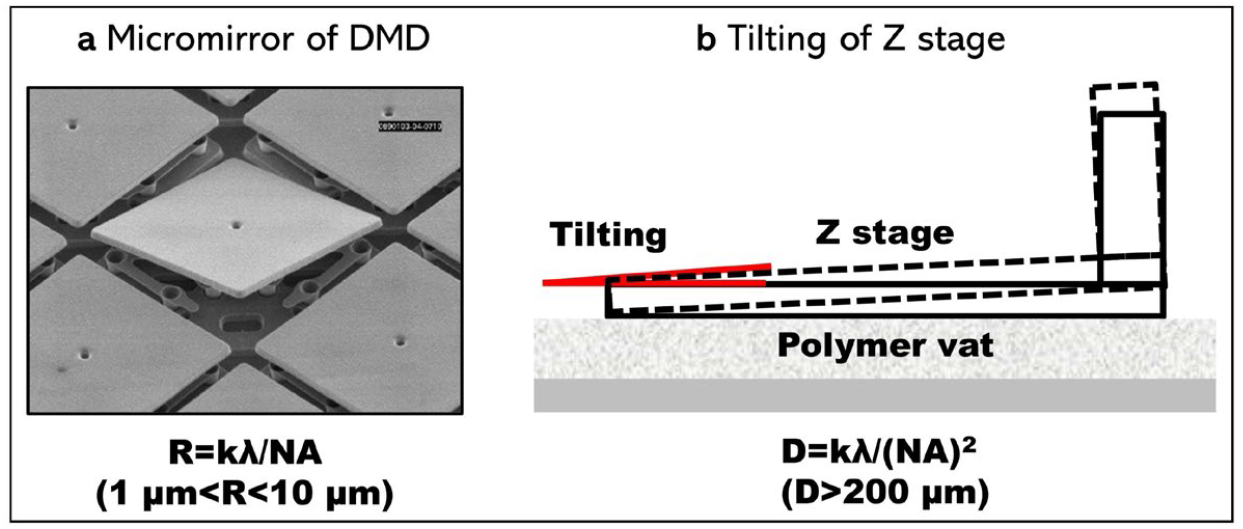
Schematic showing alignment challenges. NA is based on resolution and depth of focus.

**The modulation transfer function (MTF)** determines how much contrast in the original object (DMD) is maintained at the image plane. To resolve two adjacent pixels in DMD at the image plane. Therefore, MTF @ 50lp should be more than 0.5 for 1x optical path, and MTF @20lp should be more than 0.5 for 3x optical path.

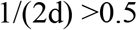

where d is the pixel size at the image plane. For 1x, d = 10.8µm. For 3x, d = 32.4 µm.

#### System design and analysis using Zemax

With these system specifications, we set the merit function in Zemax with optimization of primary aberration and the control of the magnification ratio. **Figure S3a** demonstrates the 2D layout of the optical systems. Optimization was carried out with three main field of view (FOV=0, 0.707, 1). So, the three color (Red, Green, Blue) represents the light beam from these FOVs. Zeemax was used to run system analysis, and results show that the performance for both 3x and 1x modes reach close to their diffraction limitation. *(close to ideal results)*

For 3x mode, we choose a plane-convex lens with *F=300 mm* and a tube lens with *F=200 mm*. For 1x mode, we choose a achromatic doublet with *F=200 mm* and a tube lens with *F=200 mm*. We simulated several off-shelf lens choices and found that tube lens has the better imaging performance across all the field of view due to its overall aberretion correction than standard achromats.

#### Spot diagram

Spot diagram presents the imaging performance of single dot at different field of views (FOVs). **Figure S3b** presents the spot diagram for the 3 FOVs. The three colors in this diagram presents 3 main wavelengths (0.405 µm, 0.407 µm, 0.409 µm). The RMS spot radius listed at the 3 FOVs are less than airy radius **(18.41µm)**; *airy radius represents an ideal system under diffraction limitation*. The airy radius is the black circle in **Fig. S3c**. This demonstrates that the system performance have reached its diffraction limitation.

#### Distortion diagram

The X axis of the distortion diagram is the percent distortion. The Y axis of the diagram is the FOV. Results in **Fig. S3d** show that maximum distortion is at the full field of view, which is still less than 0.1%; this meets our target specifications.

#### MTF diagram

MTF is an important index to characterize system performance. Regularly, we quantify the performance using line patterns with different spatial frequency and characterize its image contrast at the final image plane. In **Fig. S3d**, the X axis of the MTF diagram represents spatial frequency per cycle of line pattern. The Y axis of the diagram represents the modulus of the optical transfer function, which is image contrast. The three colors in this diagram represents 3 FOVs. We can see that the MTF at all FOVs is close to the ideal performance under diffraction limitation which is the dashed black lines in the figures. Even though MTF for 1× system at full field of view is still a bit less than 0.5, the system printing performance comes close to the ideal which is promising.

**Figure S3.**
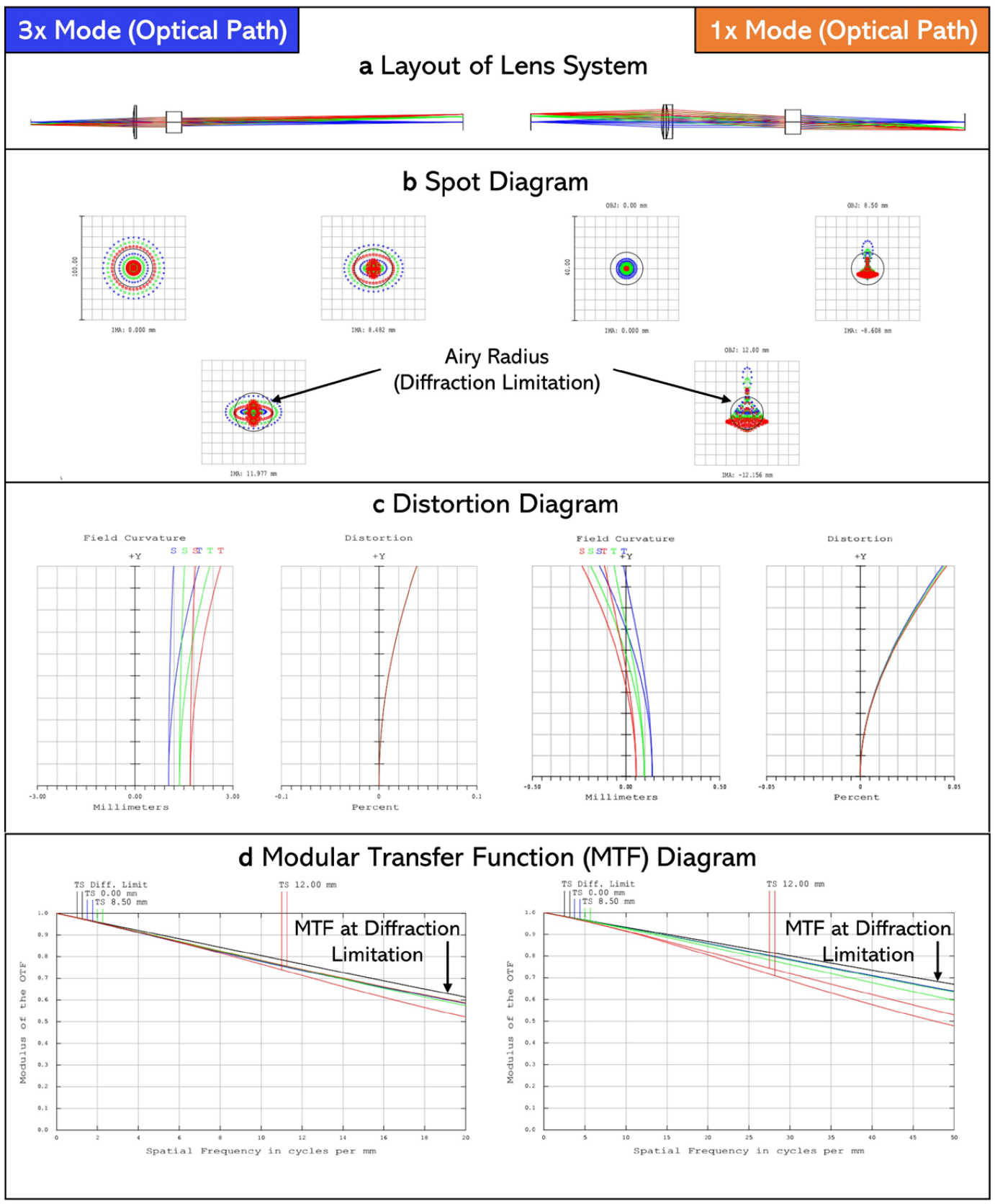
Projection lenses for both 3x and 1x modes of MPS optimized using Zemax simulations. **a** Lens layout of the system. **b** Spot diagram. **c** Field-curvature and distortion diagram. **d** MTF diagram.

### 3. MPS Printing process

#### Custom slicer

MATLAB was used to develop a custom 3D slicer to slice CAD files layer by layer into respective individual image files. The slicer was designed as a function to be run in one easy step. Once run, the user is directed through a series of prompts. This slicer can be used for all 3 modes of our printer, 1x individually, 3x individually, and 1x and 3x combined. The first prompt asks the user if they want to print with the combination mode. If yes, it then prompts the user to select each CAD file (.stl format) for 1x and 3x mode. The user is prompted after this step to select the final output folder for the images. The next prompt asks the user for the slice height for each mode in millimeters. The final step asks the user which layer of the 3x CAD the 1x features begin on. This will allow the slicer to order the images correctly in the output folder. Generally speaking, a print will have large features printed by 3x and higher resolution internal features done by 1x. Once the slicer has all the information, the slicer will individually slice each CAD file by the desired slice heights and order the images correctly based on the user input. The slicer also outputs a text file containing the flip mount mirror positions for each layer, to enable automation of the entire printing process. If the user selected no at the first step, they would be directed to use the single mode capabilities of the printer. Both 1x and 3x follow the same first 3 prompts where the user will select the desired CAD file, select the output folder for the images, and input the desired slice height. For 3x mode, the code will be complete, and the output images will be displayed for the user. At this point, the multimode and 1x individual paths have another prompt which allows for an image offset for 1x. If no offset is required, this is the final step, and the output images will be displayed for the user. The addition of a software offset to the slicer stems from perfect physical optical alignment of 1x and 3x beam paths being near impossible. Once the physical alignment is adequate, simple image processing can be used to get the alignment perfect. This process is highlighted in supplemental figures. If an image offset is needed, the user will be prompted to input an X and Y offset value in millimeters, then the output images will be displayed for the user. This is the end of the code. The current version of this slicer is version 7 and has undergone many iterations to increase the versatility of the slicer as well as meet the complex needs of the MPS printer.

#### Printer workflow

The MPS printer workflow is illustrated in **Fig. S4**. A few prepatory steps are performed before printing occurs. This includes loading the desired photoreactive material into the PDMS dish, lowering the stage to create an oxygen permeable ‘dead-zone,’ setting the mirror mount to a starting position, and inputting various printing parameters. The required parameters include laser power, the image file folder location, first layer height, first layer show time, remaining layer heights, remaining layer show times, stage velocity and acceleration, detaching distance, number of layers, and the 1x and 3x text file folder location. Now, printing is ready to begin. The DMD will be enabled and expose a UV image for the set base layer show time. Following exposure, the code will check the next layer. If the next layer is 3x, the stage will move the desired detatchment distance to allow for new material flow, then move back down the next Z layer height and expose the next layer. If the next layer is 1x, the mirrors will flip and print the 1x layer at the same Z height. Following this for either, 1x or 3x, the code will look for the next image, if there is one, the checking of the next layer process will repeat. If there are no more images, then the print will be complete.

**Figure S4.**
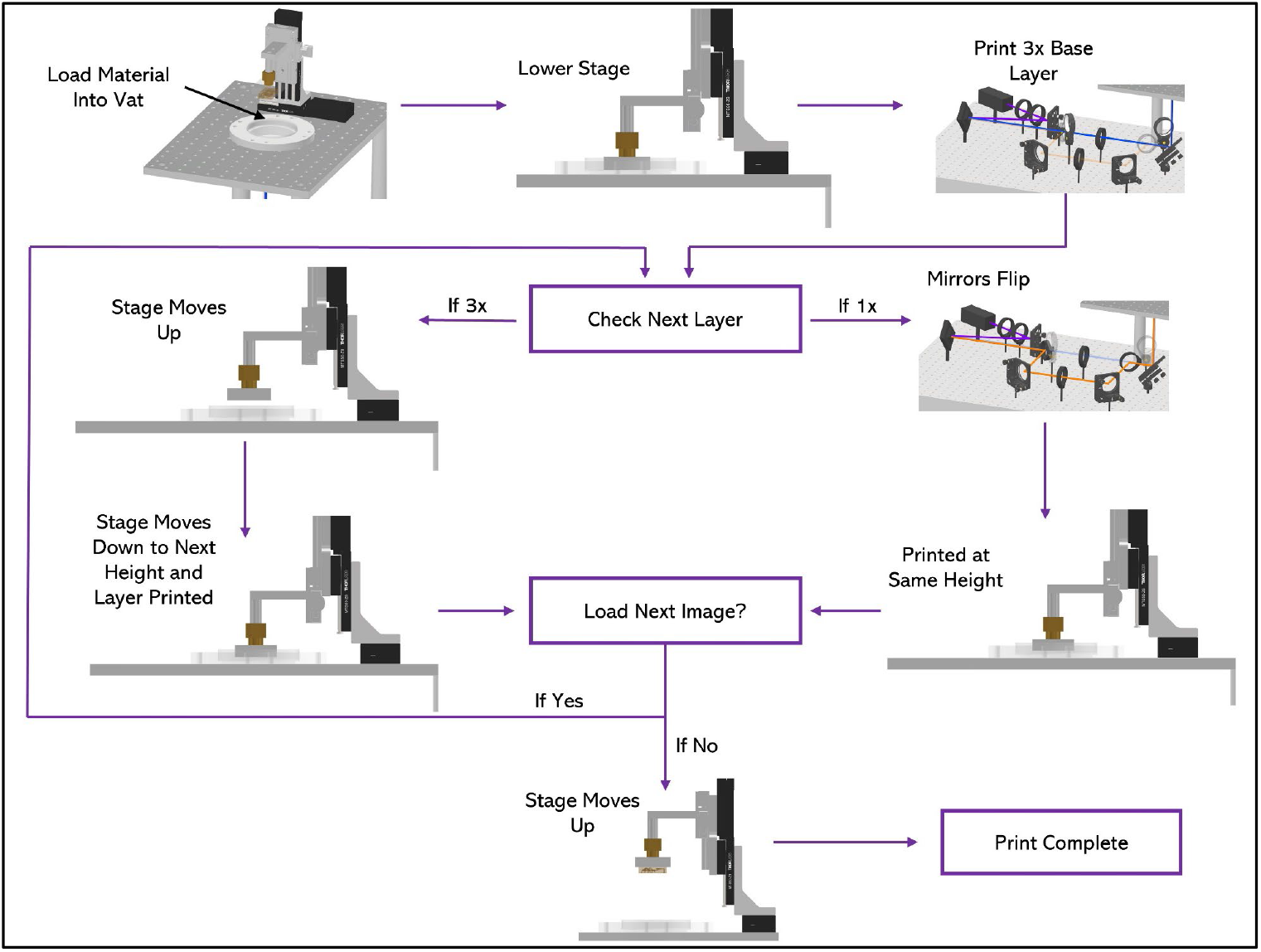
MPS process overview. Printing process start to finish (top to bottom).

### 4. Optimization of resin formulation

Two optical properties were optimized: the chosen material should crosslink or polymerize at UV or near UV wavelength and the crosslinked material should be transparent. These optical properties can be modulated by various types and concentrations of photo-initiators, and photo-absorbers. We first choose Poly(ethyleneglycol) diacrylate (PEGDA) of 250 molecular weight as the base material due to its excellent properties of swelling-resistance and impermeability to water. Then, two widely used photo-initiators were evaluated with ultraviolet (UV, 365nm) and near ultraviolet (NUV, 405nm) light sources, phenylbis(2,4,6-trimethylbenzoyl) phosphine oxide (Irgacure 819, Sigma-Aldrich) and Lithium phenyl-2,4,6-trimethylbenzoylphosphinate (LAP, synthesized in lab). Since LAP was found to be insoluble in 100% PEGDA (250mw) resin, Irgacure 819 was chosen as the photo-initiator **(Fig. S5a)**.

Next, a variety of photo-absorbers were evaluated. With light-based printing, choosing the right photo-absorber is important to obtain the highest printing resolution. Nine photo-initiators were chosen based on their wide use in the field. These included 2-isopropylthioxanthone (ITX), Sudan-I (SI), Martius Yellow (MY), 2-nitrophenyl phenyl sulfide (NPS), 2,2,6,6-tetramethylpiperidine 1-oxyl (TEMPO), Tinuvin 234 (TIN), Orange G (OG), Quinoline Yellow (QY), and Tartrazine (Tart). The photo-absorbers were then screened based on three key requirements: solubility, spectrum matching with UV and NUV wavelengths, and optical transparency OG, QY, and Tart are found to be insoluble in PEGDA (250mw) resin **(Fig. S5a)**.

TEMPO and TIN showed minimal spectral overlap with the target wavelengths of the chosen light sources, LED with peak at 365nm and Laser with peak at 405nm (**Fig. S5b)**. Insufficient spectral overlap will result in unwanted polymerization and prevent printing of voids and/or channels. Among the remaining photo-absorbers, ITX was found to be transparent while SI, MY and NPS showed yellow or orange colors **(Fig. S5c)**. ITX was the last man standing and was selected for this work. An ideal material formulation compatible with MPS exhibiting high transparency, water impermeability, and durability has been identified and was used in all outlined experiements. This included PEGDA (250mw) as the base material, Irgacure 819 as the photo-initiator, and ITX as the photo-absorber.

**Figure S5.**
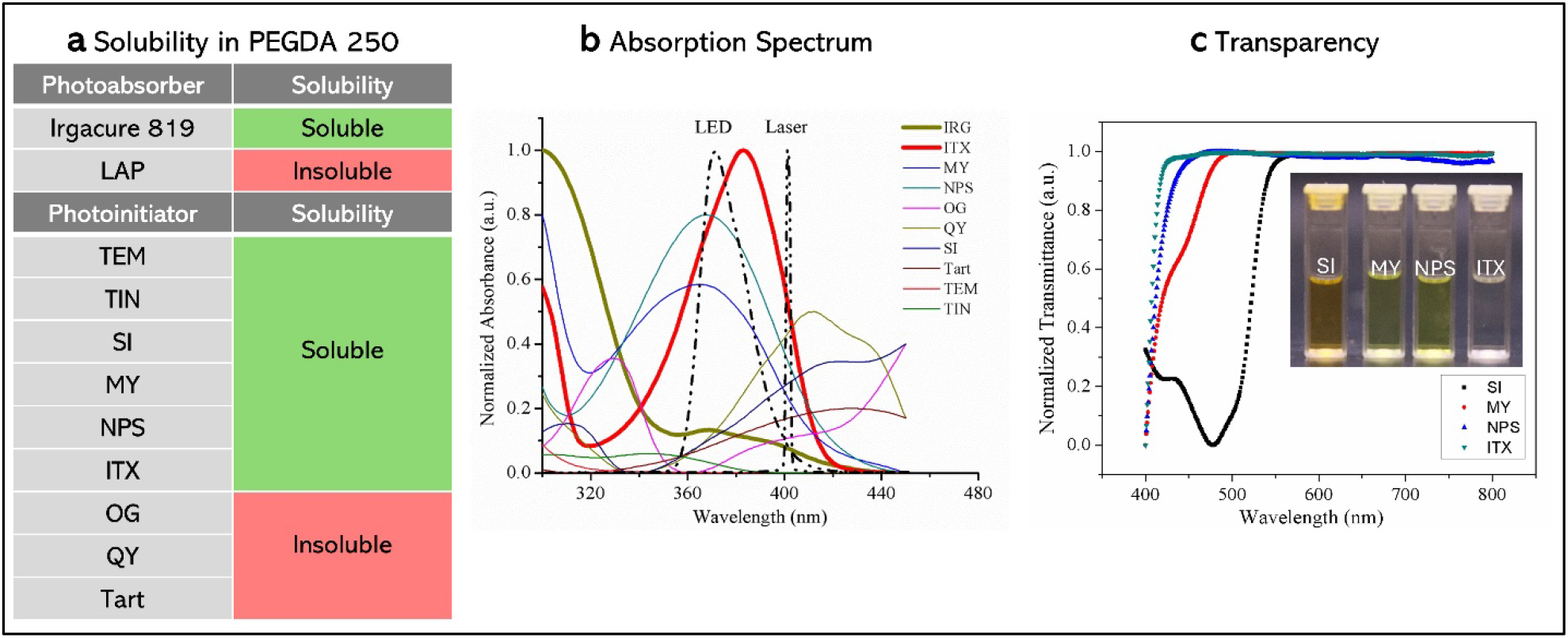
Photo-absorber screening process overview. Nine photo-absorbers selected from their wide use in the field and narrowed down based on three key requirements for this work. **a** Solubility in PEGDA 250 (base material) was first evaluated, leaving six photo-absorbers. **b** The absorption spectrum of the photo-absorbers was evaluated comparing them to that of a UV LED and laser light source, leaving four absorbers. **c** The remaining photo-absorbers were tested for transparency, and ITX was identified as the best candidate, fitting all of the evaluation criteria.

### 5. Computation fluid dynamics (CFD)

#### CFD

Computational fluid dynamics analysis was completed on the mixers to determine real world mixing efficiency.

#### Governing equations

The concentration field for the red and the blue dye in the domain is calculated by solving two independent convection-diffusion equations at steady-state conditions.

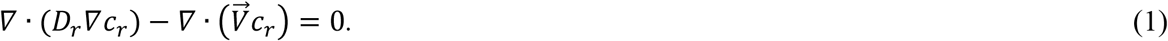

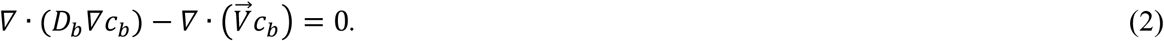

Where *D*_*r*_ and *c*_*r*_ are the diffusivity and the concentration of the red dye, and *D*_*b*_ and *c*_*b*_ are the diffusivity and the concentration of the blue dye, respectively. The independence of Eq. 1 and Eq. 2 is a valid assumption at low solute concentrations where the diffusivity of the red dye is independent of the concentration of the blue dye and vice versa. The velocity field 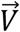 in Eq. 1 and Eq. 2 is computed by solving the incompressible continuity and Navier-Stokes equations at steady-state conditions.

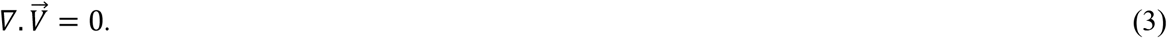

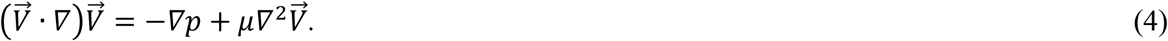

Where P is the pressure field and *µ* is the dynamic viscosity of the solvent fluid. Here, we assumed that the velocity field is independent of the concentration field. This is valid when the concentration of the solute is small; thus, its effect on the density and the viscosity of the fluid is negligible.

#### Determining the diffusion coefficients

The diffusivity of a solute in a solvent can be estimated by the Stokes-Einstein relation.

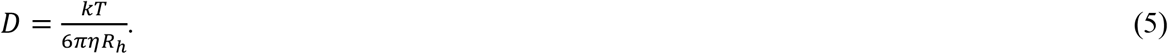

Where *K* is the Boltzmann constant, *T* is the temperature of the solution, *η* is the kinematic viscosity of the solvent, and *R*_*h*_ is the hydrodynamic radius of the solute molecules. The hydrodynamic radius for FITC-Dextran *M*_*w*_ = 150 × 10^3^ g/mol (blue dye) is R_h,b_ ≈ 85 Å according to the manufacturer Sigma-Aldrich and the hydrodynamic radius for RITC-Dextran M_w_ = 10 × 10^3^ g/mol (red dye) is R_h,r_ ≈ 23.6 Å according to the manufacturer TdB labs. These values are close to those obtained by empirical relation R_h_ = 0.488*M*_*W*_^0.437^ for dextran [1]. Using Eq. 5 for T = 20 °C, and 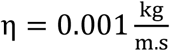 for water; we determined D_b_ = 2.52 × 10^−11^ m^2^/s and, D_r_ = 9.09 × 10^−11^ m^2^/s. The calculated diffusion coefficients are consistent with the empirical relation D = 7.69 × 10^−5^M_W_^−0.48^ [2, 3].

#### Simulation setup

Water enters from each inlet at the mass flow rate 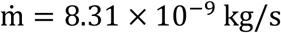 and atmospheric pressure is set at the outlet. The no-slip boundary condition is imposed on all the channel walls. We defined two scalar transport equations in Fluent corresponding to Eq. 1 and Eq. 2. SIMPLE scheme was chosen to solve the pressure-velocity field [4]. The Green-Gauss node-based gradient evaluation with a second-order accuracy is used for spatial discretization of the momentum equation and scalar transport equations [5].

#### Case No. 1 - Fixed solid wall fins

The geometry of the domain is shown in **Fig. S6**. The maximum velocity in the channel is V_max_ = 0.000427 m/s, and the characteristic length of the channel is L = 0.16 mm; thus, the Reynolds number 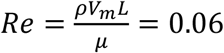, which is well within the laminar region.

**Figure S6.**
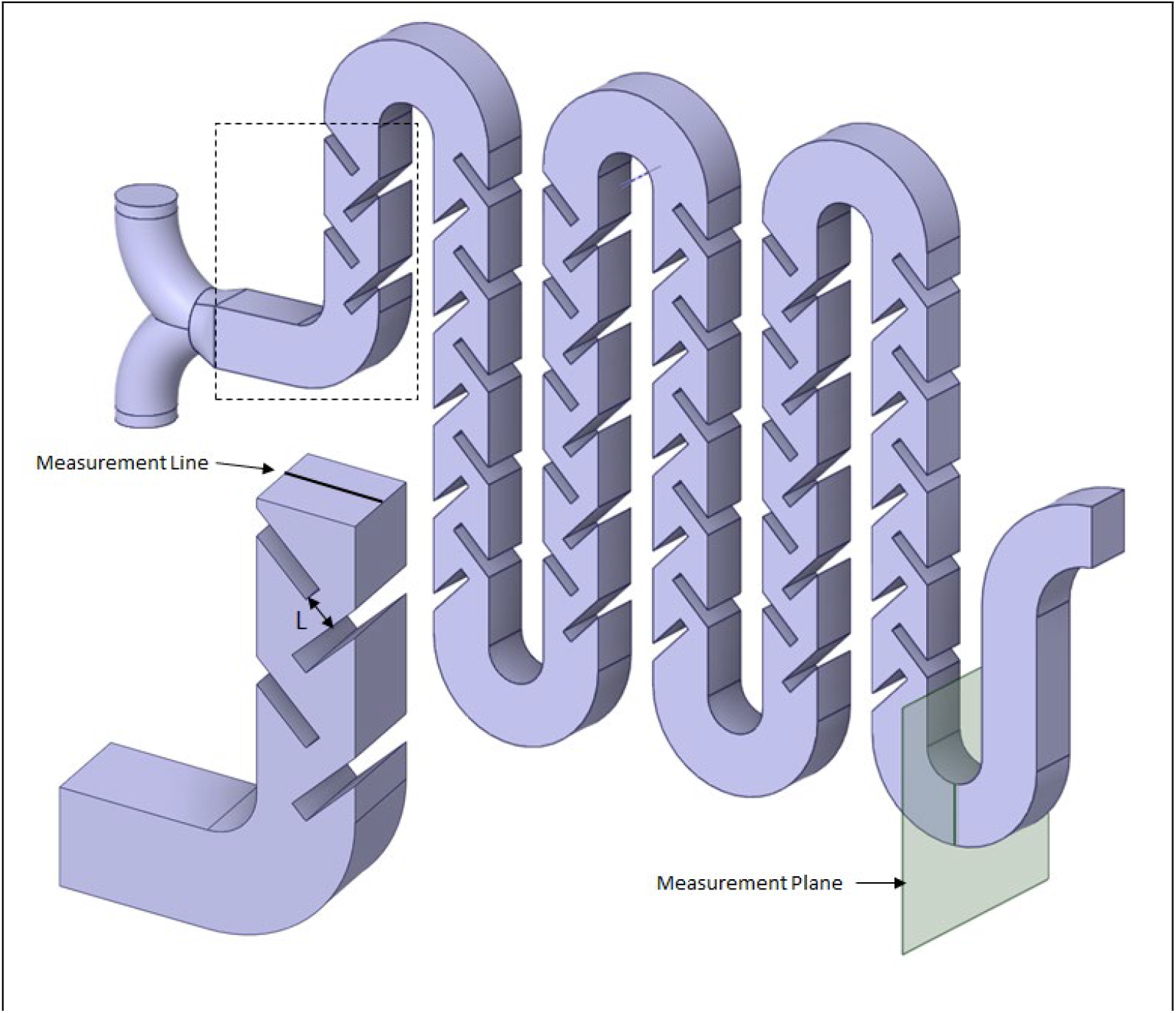
Microfluidic mixer geometry number 1.

We performed a mesh independence test to find a compromise between computation accuracy and cost. We conducted the mesh test on the section of the geometry shown in the bottom left corner of **Fig. S6**. We selected the number of mesh cells, n, per characteristic length of channel L, as the reference. We conducted five simulations for *n* = 10, 15, 20, 25, 30 which correspond to the total number of tetrahedral cells N = 769654, 3037784, 6105443, 15226105, 21375077. We ran the simulations until the scaled residuals for the scalar equations (corresponding to Eq. 1 and Eq. 2) converged. For Eq. 1, the residual converged at 3.6 × 10^−8^, 7.5 × 10^−14^, 1.1 × 10^−16^, 5.2 × 10^−14^, 1.5 × 10^−15^, respectively as we increased *n*. For Eq. 2, the residual converged at 4.8 × 10^−7^, 1.4 × 10^−7^, 7.7 × 10^−8^, 2.2 × 10^−8^, 2.5 × 10^−8^, respectively.

**Figure S7.**
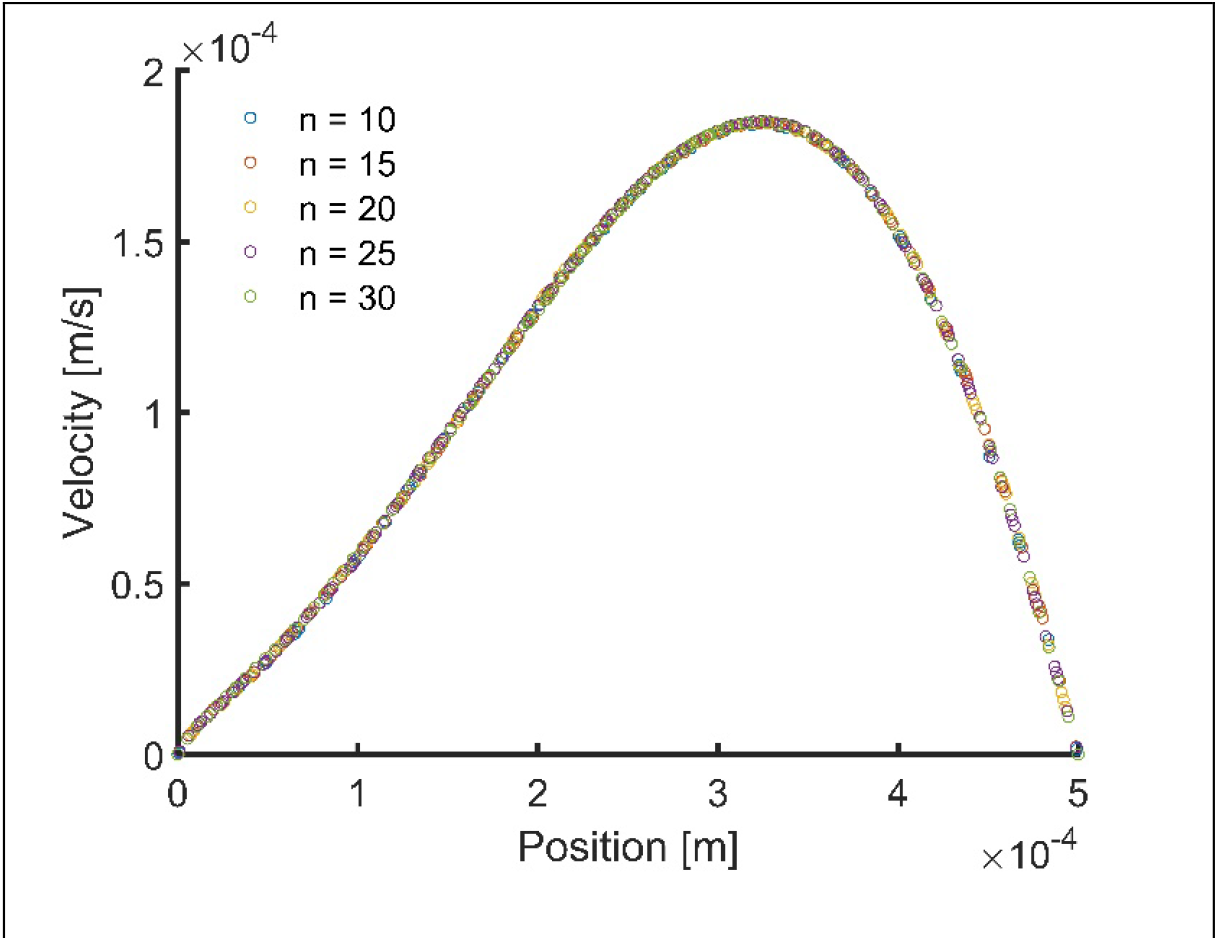
Flow velocity on the centerline of the outlet.

**Figure S8.**
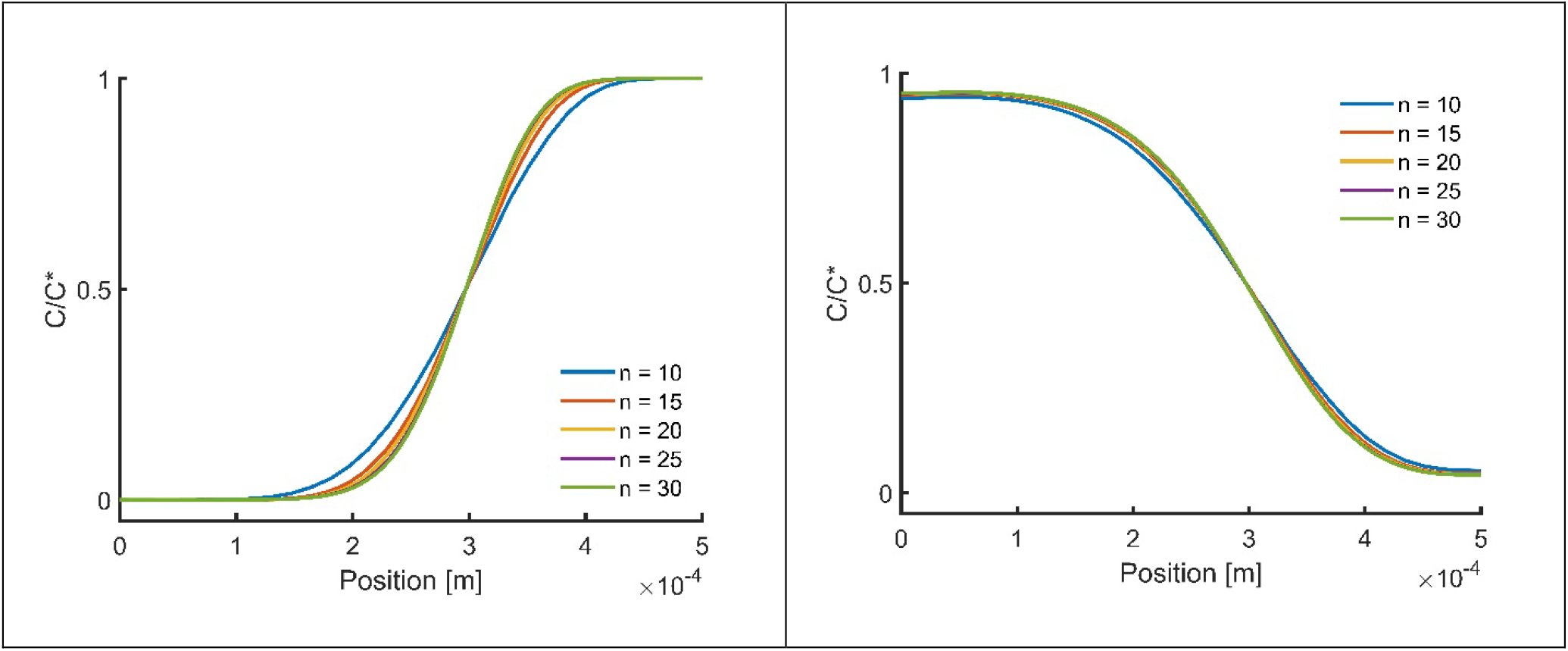
Concentration of the red (left) and the blue (right) dyes on the centerline of the outlet.

**Fig. S7** shows the velocity profile on the centerline of the outlet (shown in **Fig. S6**) for the different values of n. The solution does not change with the value of n, and thus is independent from the mesh size. **Fig. S8** shows the variation in the concentration of the red and the blue dye on the centerline of the outlet. The concentration field of the red dye is still well in the mesh size-dependent region, whereas the concentration of the blue dyes is relatively less sensitive to the mesh size. As our final goal of this setup is to determine the degree of mixing between the blue and the red dye, we calculated a mixing efficiency index *ME* at the outlet using the results of the five simulations shown in **Fig. S9**. We will express the formula for the mixing efficiency in the next section. *ME* decreases from 19.48% to 18.98*%* when n increases from 20 to 25. This is 2.56% change, which becomes less than 1% between n = 25 and n = 30. We chose n = 20 for the primary simulation, corresponding to 172647199 mesh cells.

**Figure S9.**
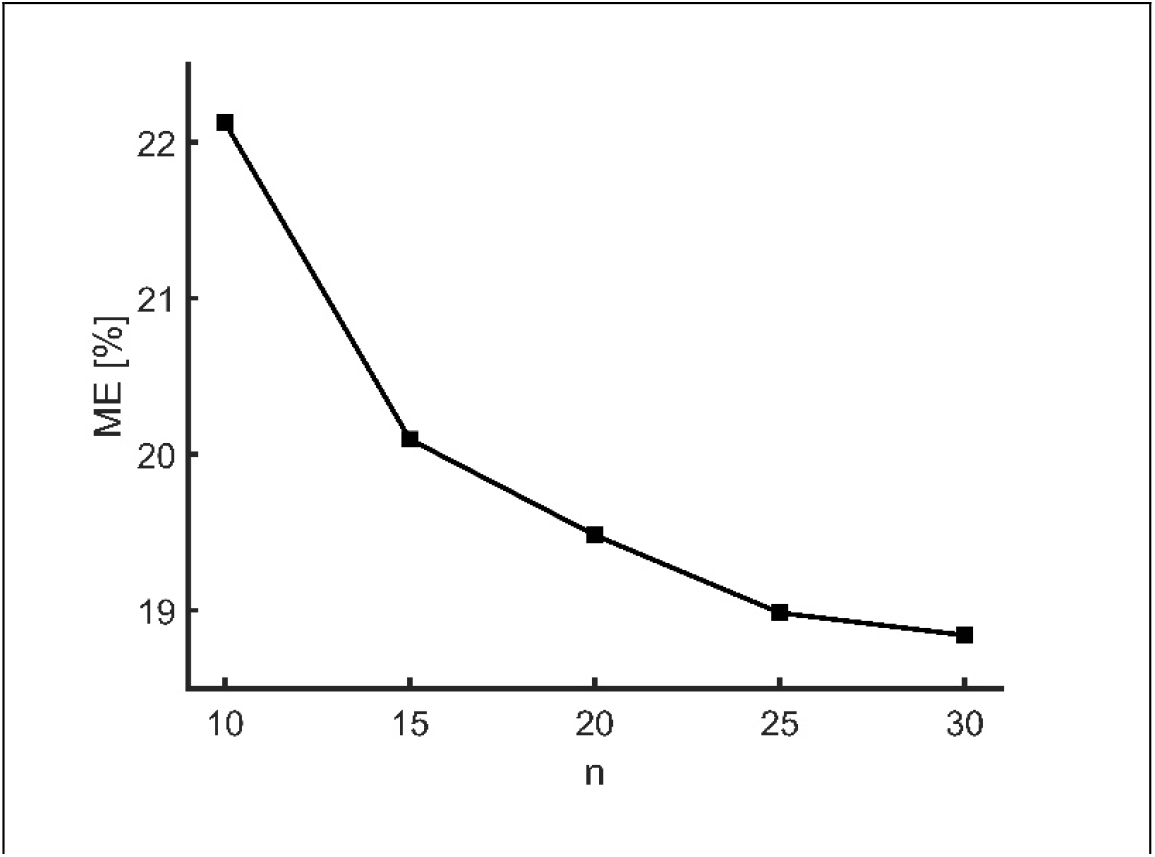
Mixing efficiency (ME) versus n.

**Case No. 2 – 3D spiral fins. Fig. S10** shows the third geometry studied in this work. We performed a mesh dependency test on the entire domain using three mesh element lengths of *l* = 1.9 × 10^−5^ *m, l* = 1.2 × 10^−5^ *mm* which correspond to 8149834 and 36412425 cells, respectively. In the first trial, the residuals corresponding to equation 1 and equation 2 converged at 1.32 × 10^−9^ and 3.36 × 10^−13^, respectively. For the second trial the residuals converged at 1.47 × 10^−10^ and 8.48 × 10^−12^, respectively.

**Figure S10.**
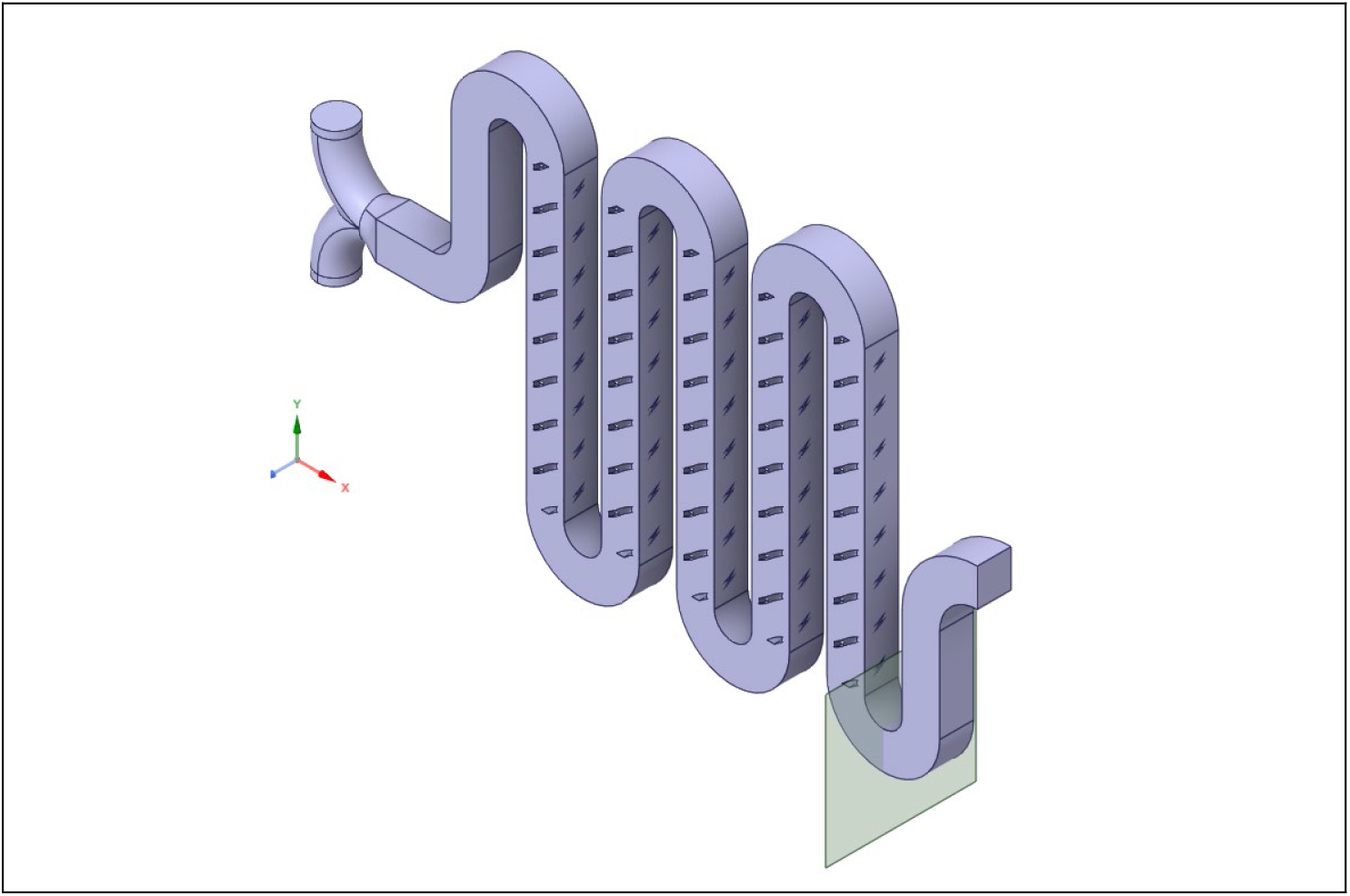
Microfluidic mixer geometry number 2.

The velocity profile on the centerline of the measurement plane is shown in **Fig. S11**. The changes in the velocity are negligible as the result of reducing the mesh size and we conclude that the velocity field is independent of the mesh size. **Fig. S12** shows the concentration of the red and blue dye on the line of interest. The concentration of the red (**Fig. S12a**) and the blue dye (**Fig. S12b**) on the centerline is sensitive to the mesh size. Although the concentration field is sensitive to the mesh size, the differences are very small. The micing efficiency changes from 99.6% to 99.18% as the mesh size decreases. This is less than 1% change and we are satisfied with the current results and conclude that no further mesh refinement is necessary.

**Figure S11.**
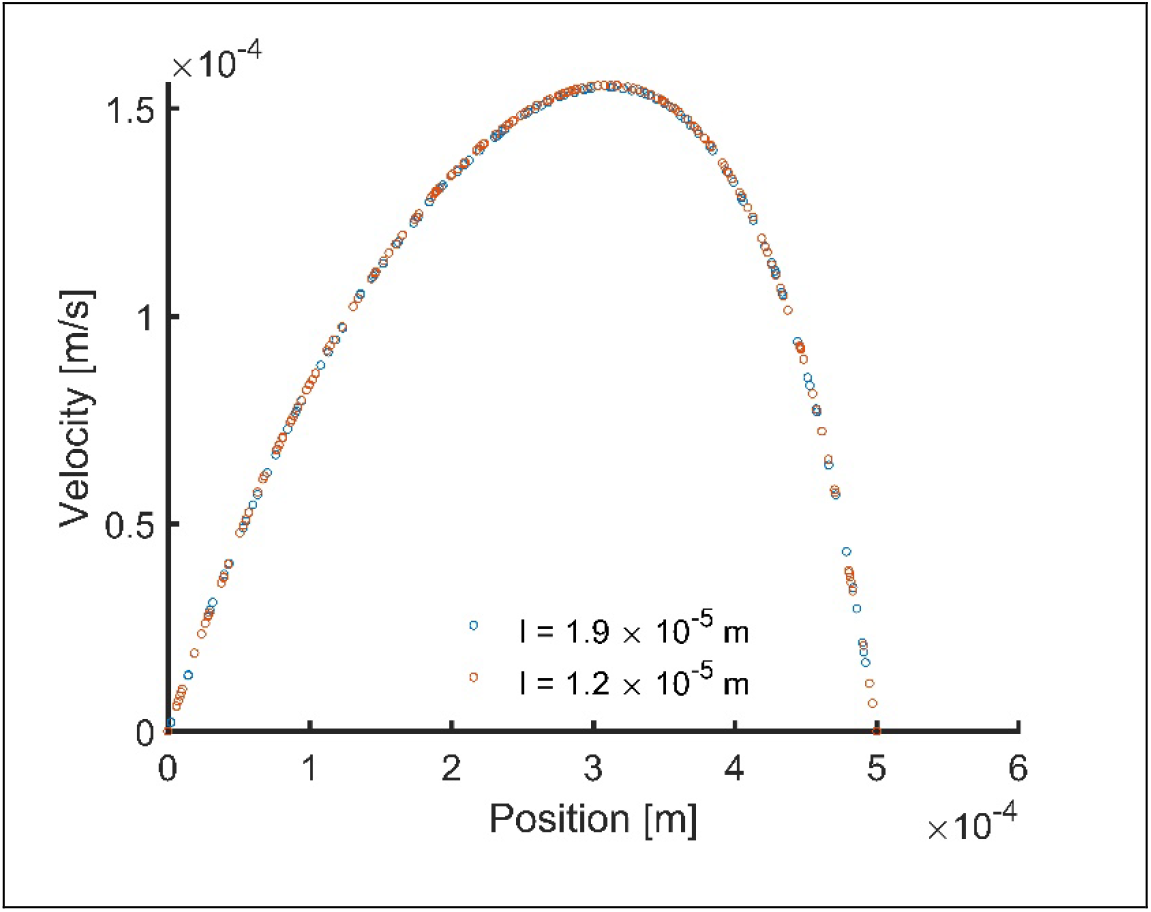
Flow velocity on the centerline of the measurement plane.

**Figure S12.**
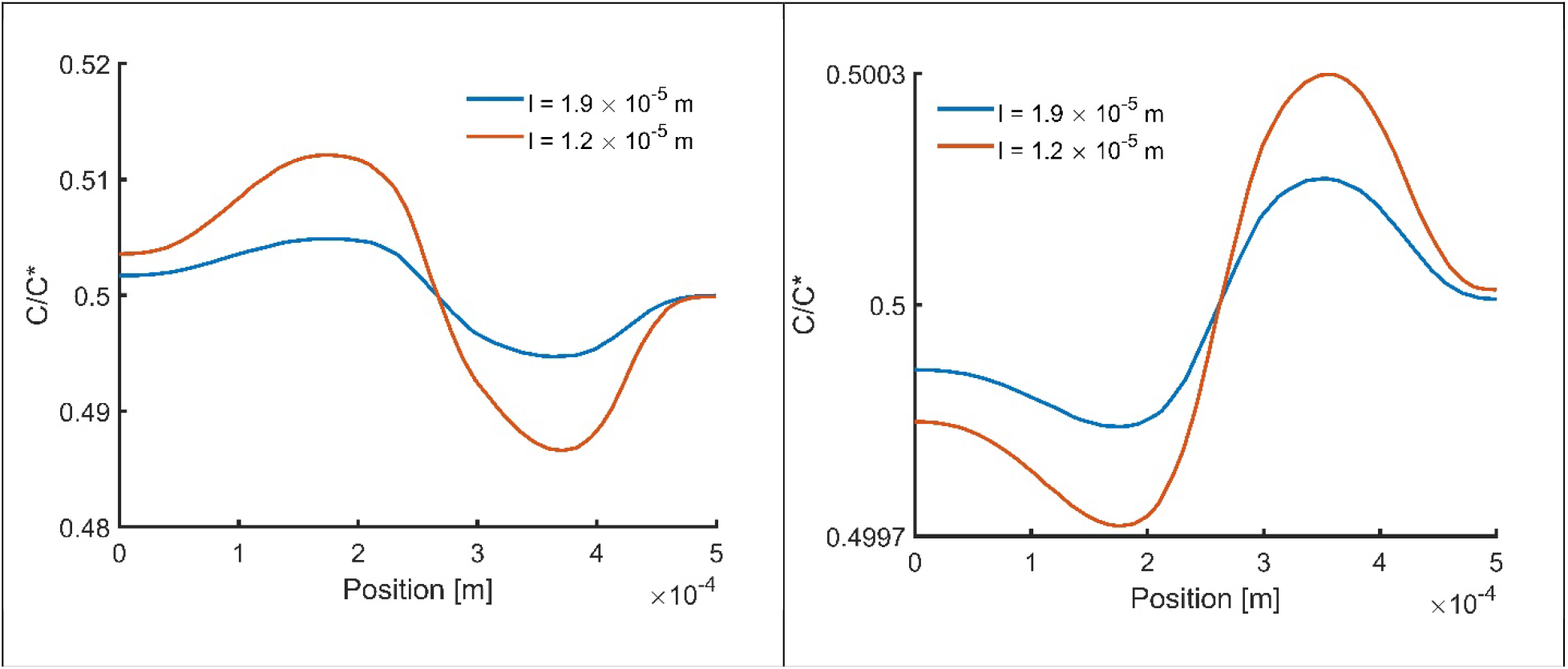
Concentration of the red (left) and the blue (right) dyes on the centerline of the outlet.

**Case No. 3 – Herringbone pattern fins. Fig. S13** shows the second geometry we studied in this work. The maximum velocity in the channel is V_max_ = 0.00019 m/s. The characteristic length of the channel is L = 0.35 mm which gives 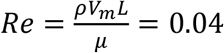. We conducted the mesh dependence test on the entire domain. For the first trial, we chose the length of a mesh cell *l* = 1.9 × 10^−5^ *mm* which corresponds to 9235333 tetrahedral cells on the entire domain. In each trial we run the simulation until the residuals corresponding to Eq. 1 and Eq. 1 converge. For the first trial the residuals converge at 1.44 × 10^−9^ and 6.31 × 10^−8^ for Eq. 1 and Eq. 2 respectively. For the second trial we set *l* = 1.2 × 10^−5^ *mm* which results in 36433691 total mesh cells. The residuals corresponding to Eq. 1 and Eq. 2 converge at 1.62 × 10^−9^ and 1.42 × 10^−8^, respectively. **Fig. S14** shows the velocity profile on the centerline of the measurement plane shown in **Fig. S13**. The change in the velocity profile is negligible as we decreased the mesh size and thus, we conclude that the velocity profile is in the mesh independent region. The effect of mesh size on the concentration profile on the centerline at the measurement plane is depicted in **Fig. S15**. Finally, we calculate the *ME* = 68.38*%* and *ME* = 66.2*%* corresponding to *ll* = 1.9 × 10^−5^ *m* and 1.2 × 10^−5^ *m*, respectively. This change is 3.19*%* and we expect it to become smaller with decreasing the mesh size and thus we conclude that the results for 1.2 × 10^−5^ *m* are accurate enough for our purpose.

**Figure S13.**
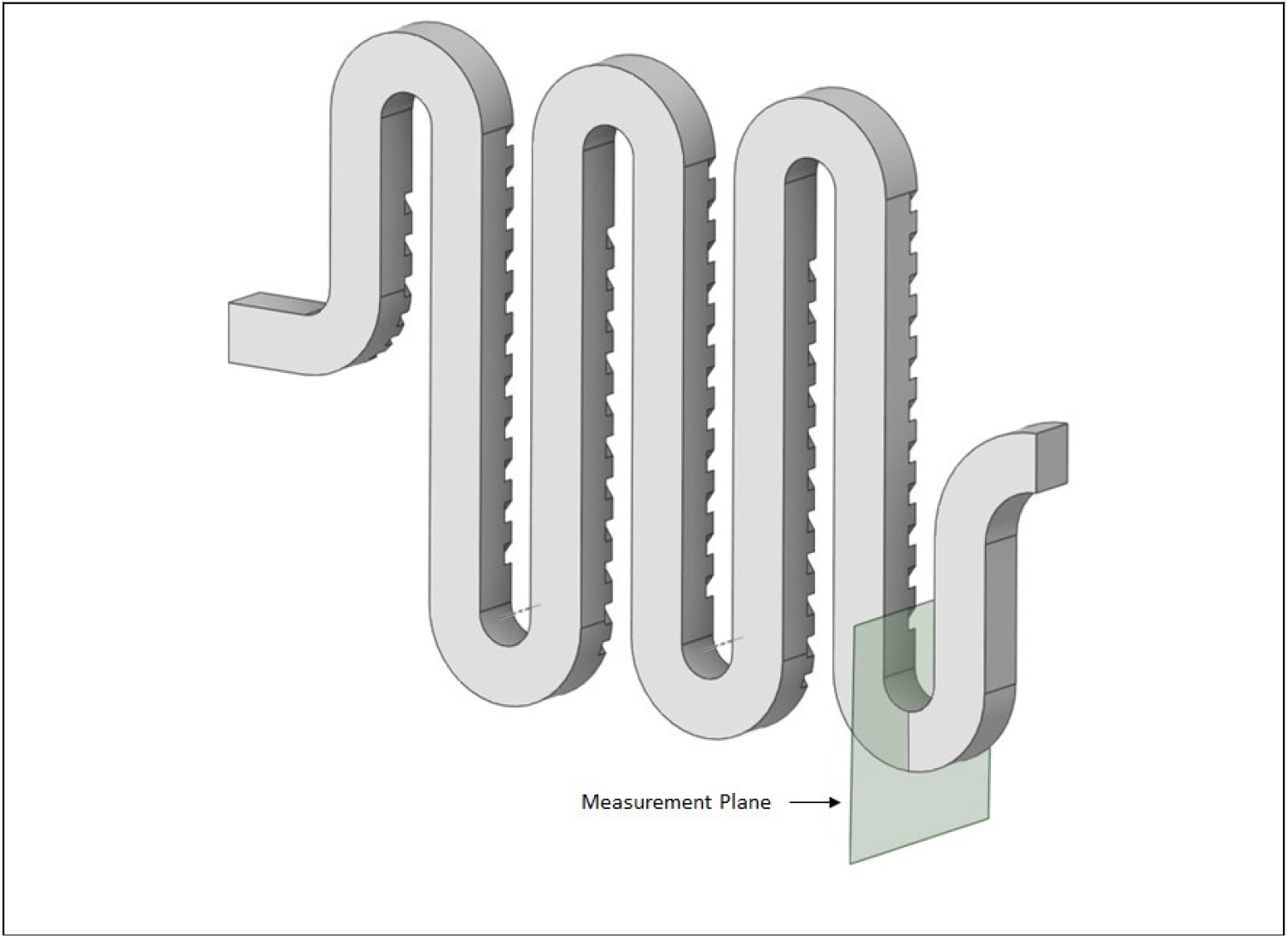
Microfluidic mixer geometry number 3.

**Figure S14.**
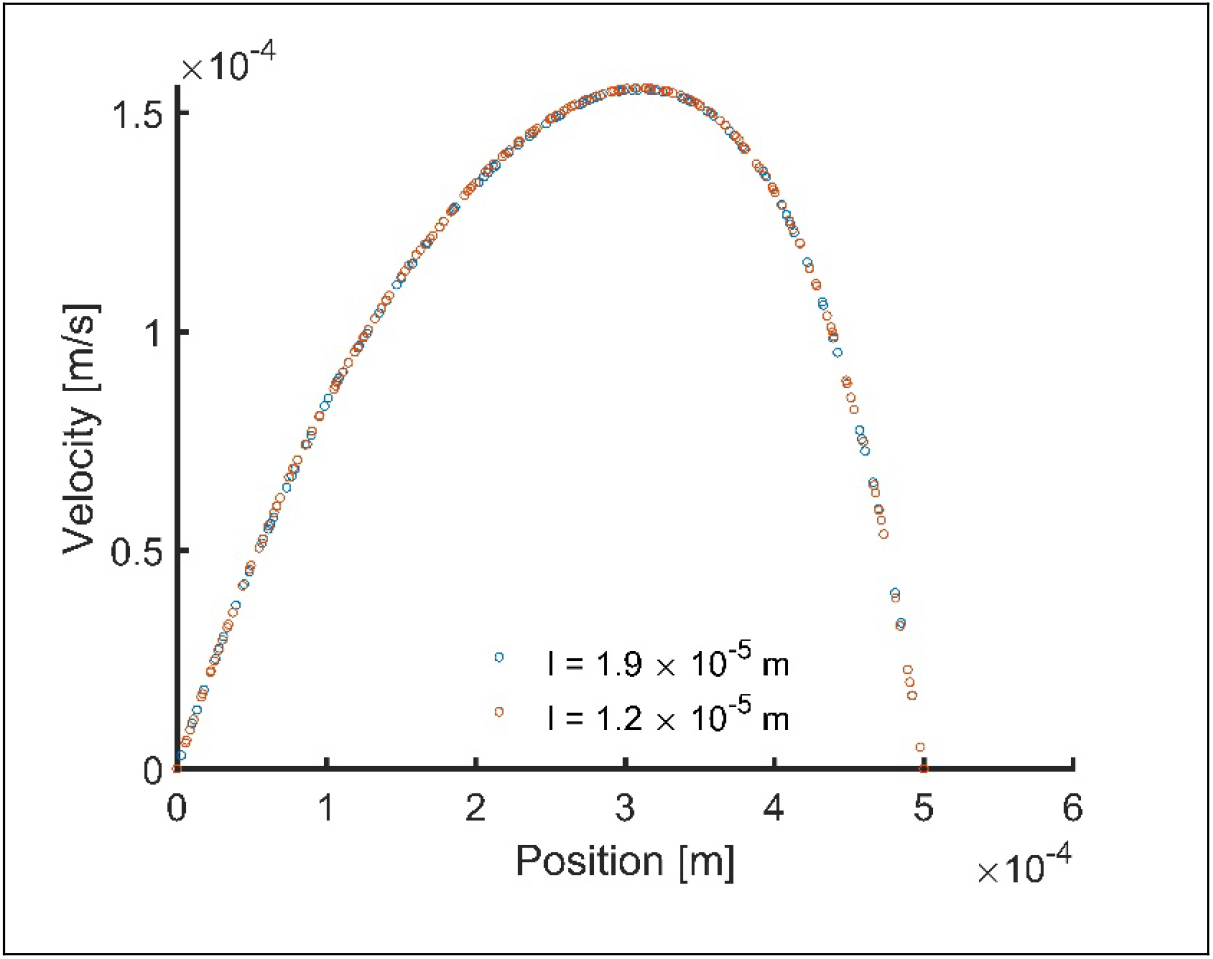
Flow velocity on the centerline of the measurement plane.

**Figure S15.**
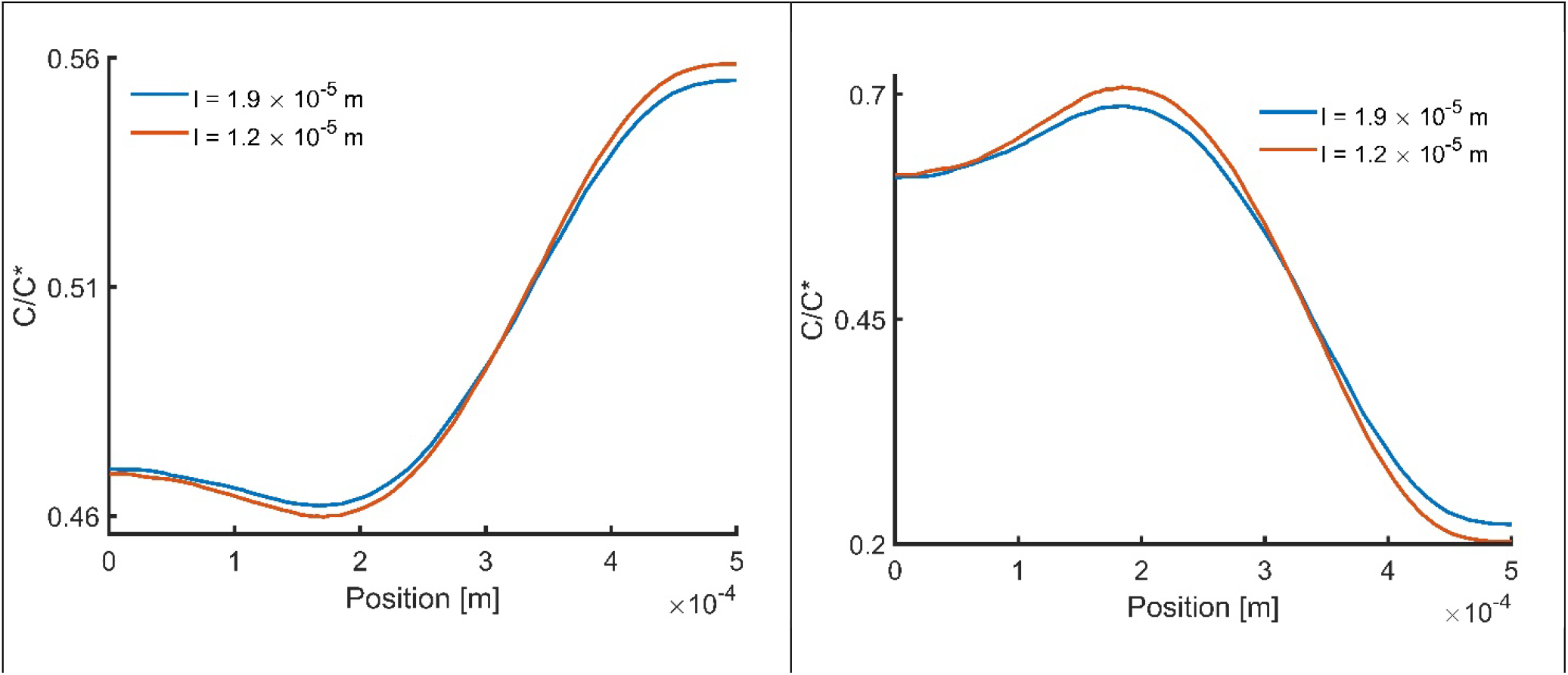
Concentration of the red (left) and the blue (right) dyes on the centerline of the outlet.

#### Calculation of mixing efficiency

Mixing efficiency of one species in a homogenous medium at any cross section *A* of a channel can be calculated by the following relation.[6, 7]

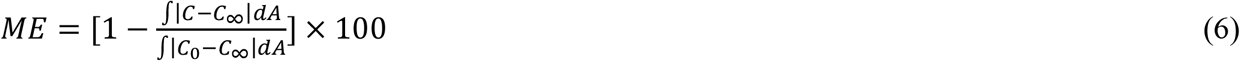

Here we are interested in calculating the mixing efficiency of two species with each other and therefore we define a mixing ratio between the red and the blue dye as follows.

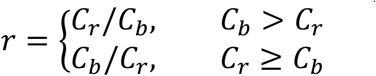

This relation ensures that 0 *≤ r ≤* 1 where *rr* = 0 and *r* = 1 correspond to no mixing and well-mixed conditions at any given point in the domain. Now we can use Eq. 6 to find the average value of *r* for any cross section of the channel which we call the mixing efficiency between the red and the blue dye.

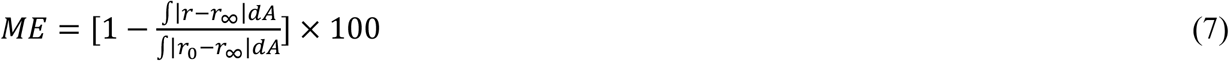

*ME* = 0*%* at the inlet and it will increase downstream of the channel with a maximum value of 100%.

### 6. Microfluidic mixers

**Figure S16.**
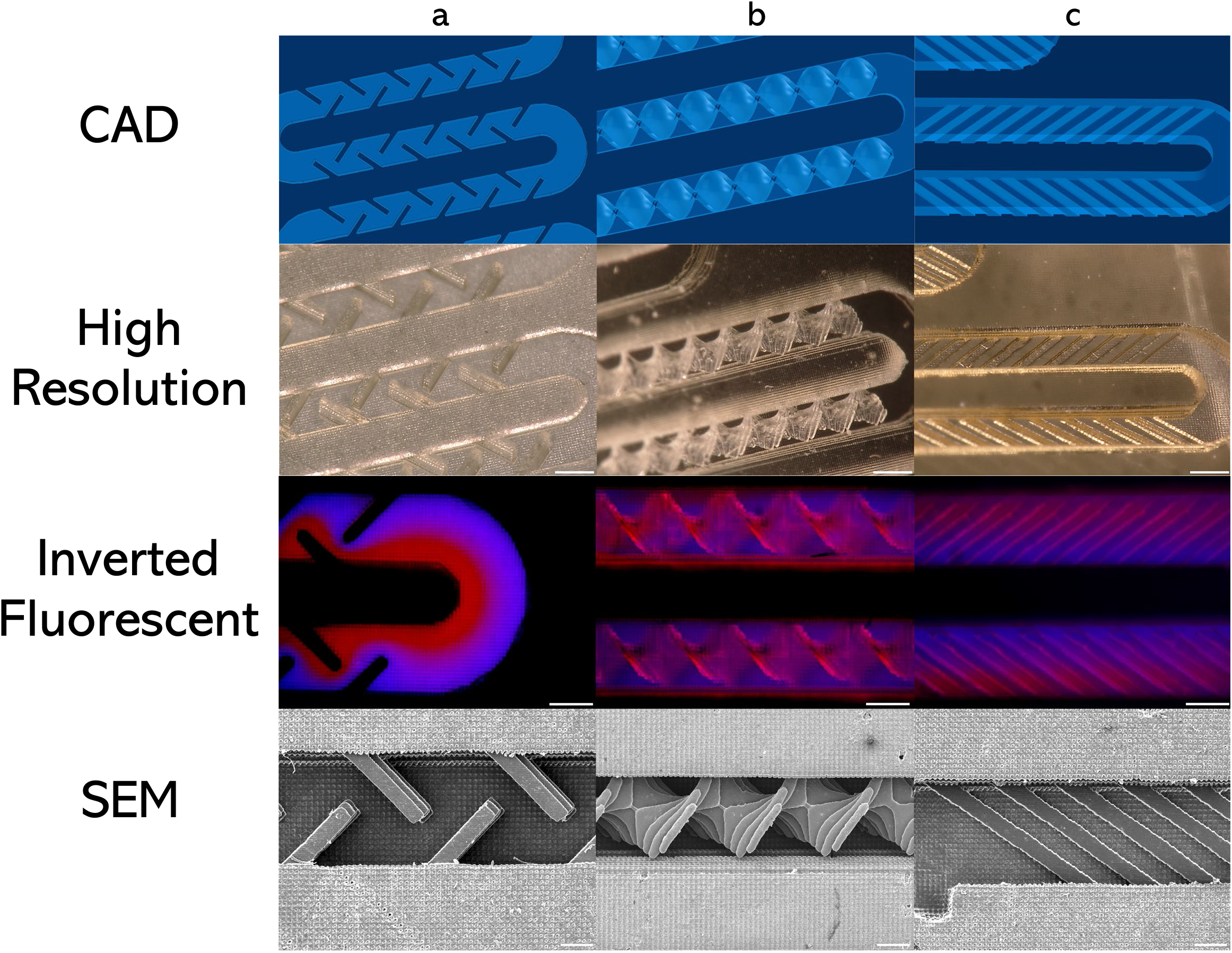
Additional images for each mixer design including **a** fixed solid wall **b** 3D spiral and **c** herringbone design. Respective isometric CAD, HIROX, inverted fluorescent, and SEM images are shown. Scale bars are 500µm (High-res), 500µm (Fluorescent), and 200µm (SEM).

**Figure S17.**
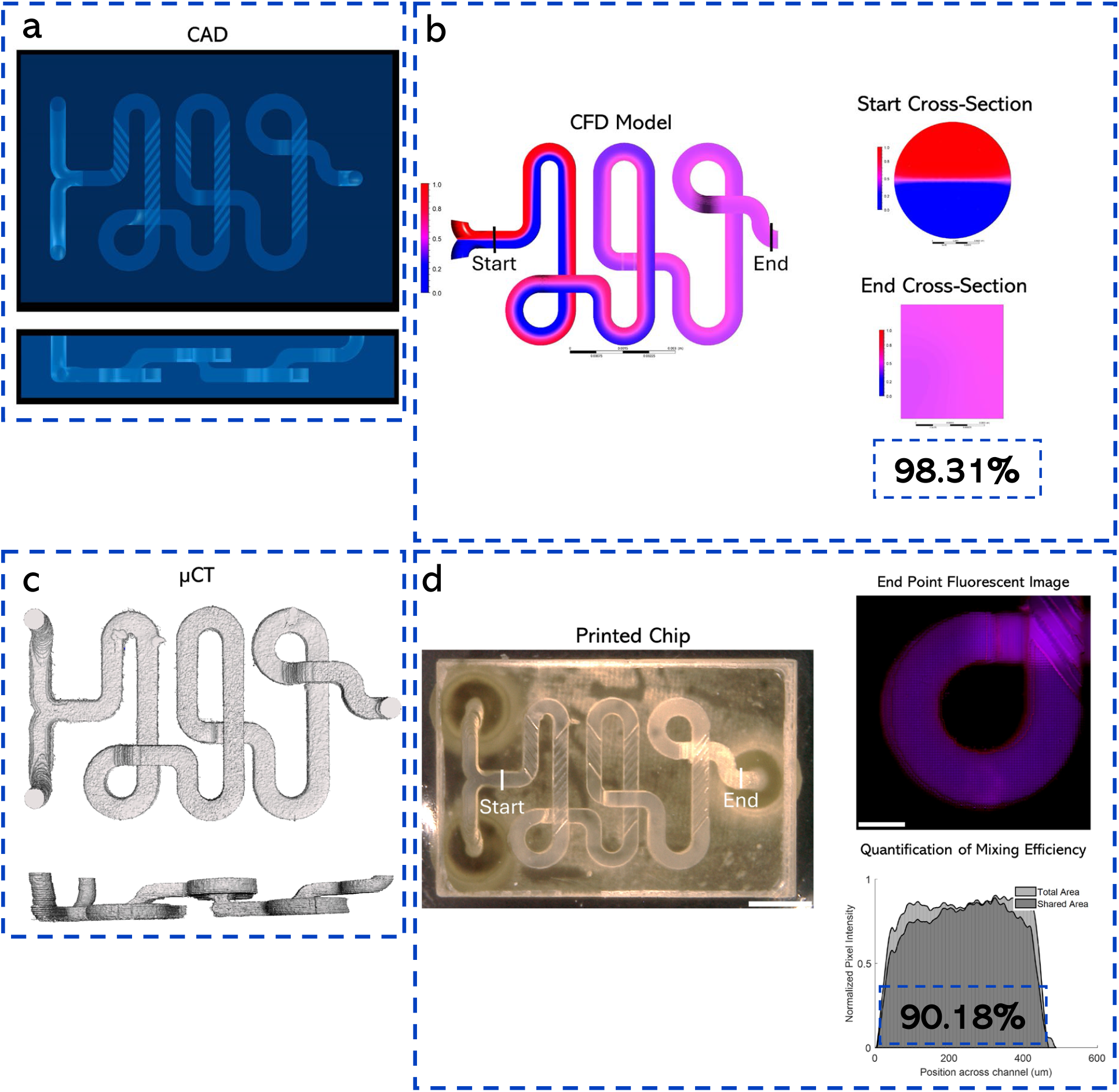
**a** CAD of complex 3D microfluidic mixer including top and side view. Design includes herringbone pattern on bottom of channels on two planes in 3D space. **b** CFD results include top view and start/end cross section views. **c** microCT reconstruction of microfluidic mixer with same model views. **d** Printed result, top view and end point fluorescent image with mixing efficiency graph. Scale bars are 2.5mm and 500µm.

**Figure S18.**
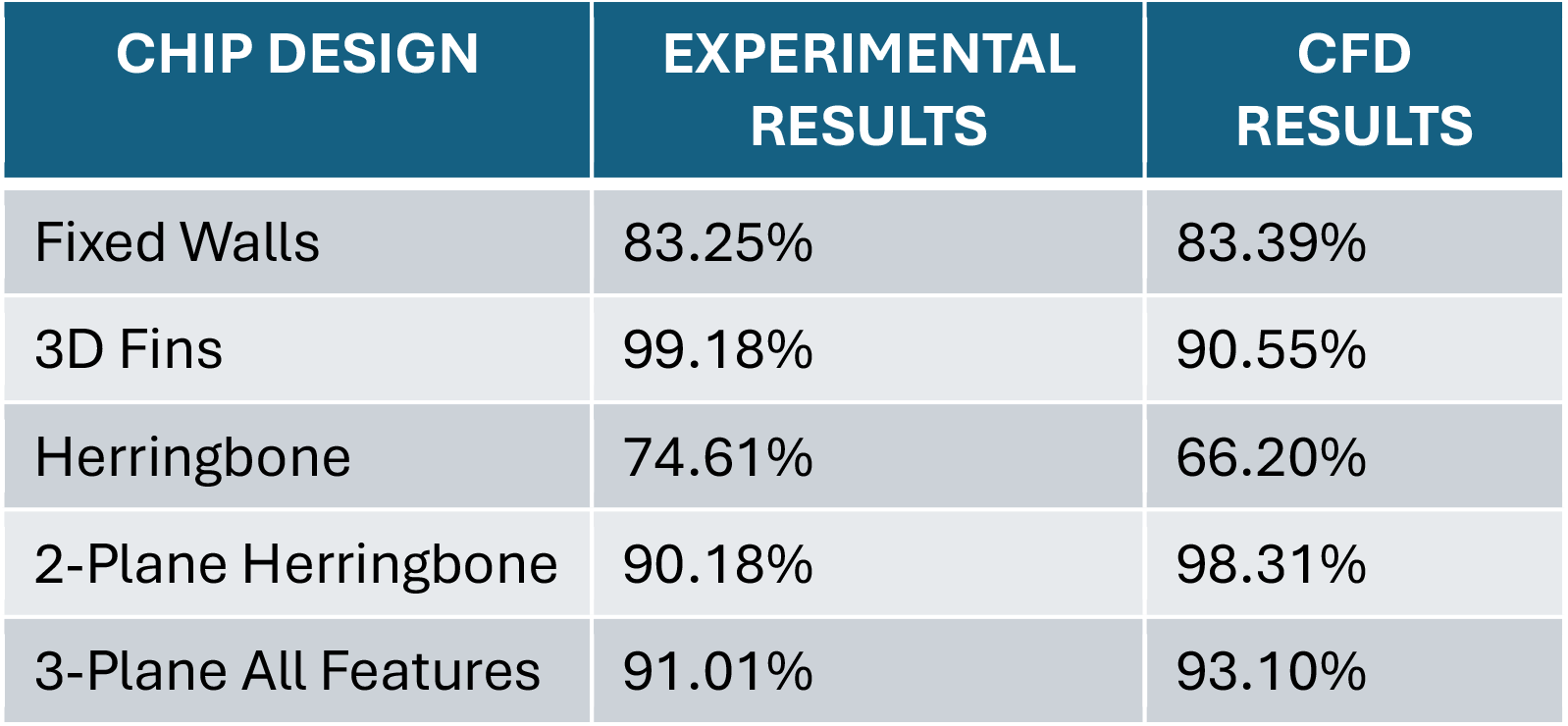
Mixing efficiency summary table.

**Figure S19.**
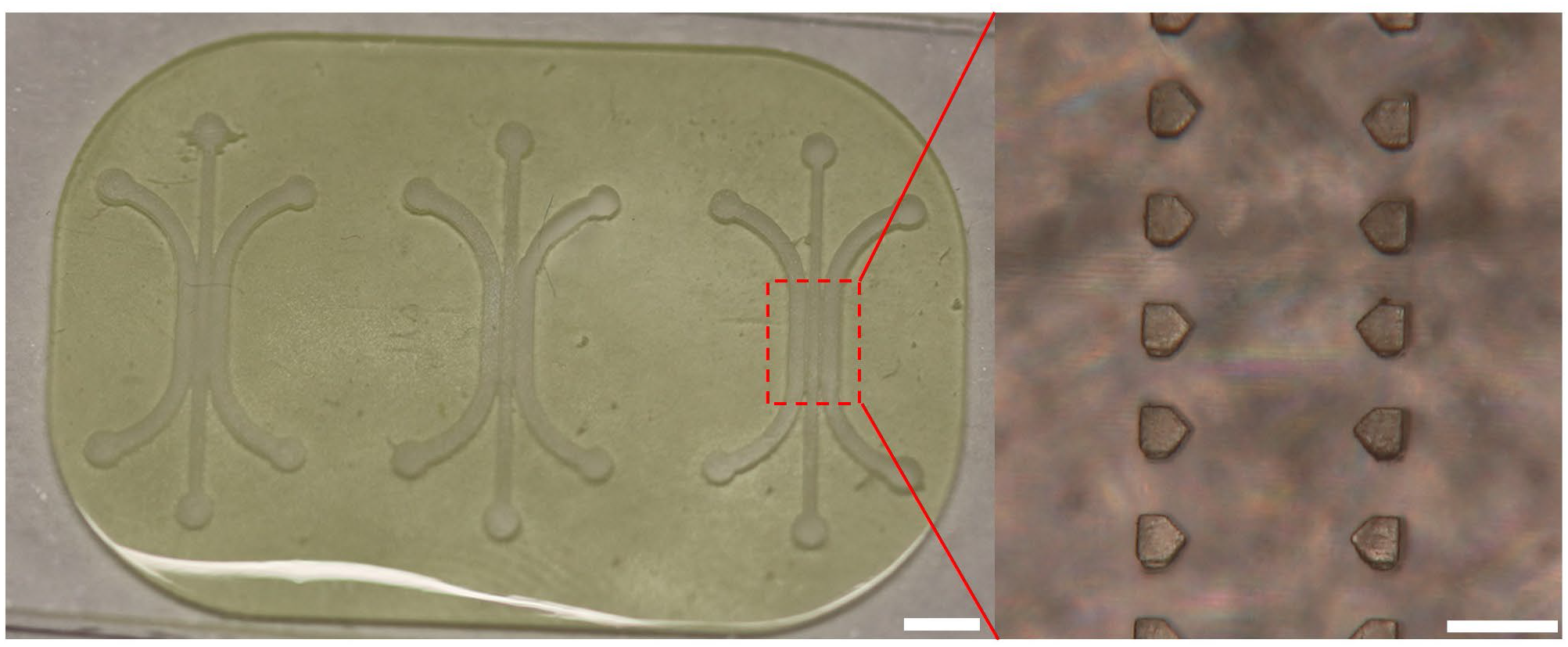
Printed three-channel cell communication chip. Zoom in of 1x features inside each chip. Scale bars are 2mm and 200µm.

